# Lipid-Mediated Control of ER Function During Apoptosis

**DOI:** 10.1101/2025.12.01.691522

**Authors:** José Luis Garrido-Huarte, Lucía López-Gil, Josep Fita-Torró, Miguel Heredia, Amparo Pascual-Ahuir, Markus Proft

**Affiliations:** Department of Metabolism, Inflammation and Aging, Instituto de Biomedicina de Valencia IBV-CSIC, 46010 Valencia, Spain; Valencia Biomedical Research Foundation, Centro de Investigación Príncipe Felipe (CIPF) - Associated Unit to the Instituto de Biomedicina de Valencia IBV-CSIC, Valencia, Spain; 2Grupo de Ingeniería Biomolecular y Biosensores, Centro de Investigación e Innovación en Bioingeniería Ci2B, Universitat Politècnica de València, Ciudad Politécnica de la Innovación, Edificio 8B, Camino de Vera s/n, 46022 Valencia, Spain; Universitat Politècnica de València, Ph.D. program in Biotechnology, Camino de Vera s/n, 46022 Valencia, Spain

**Keywords:** trans-2-Hexadecenal, Endoplasmic Reticulum Stress, Unfolded Protein Response, Fmp52/Tip30, Kar2/BiP lipidation, Lipid-Induced Apoptosis

## Abstract

Lipid-derived metabolites act as potent regulators of cell fate by disrupting organelle homeostasis, yet the mechanisms by which they trigger apoptosis remain incompletely understood. Here, we investigate how the sphingolipid-derived aldehyde trans-2-hexadecenal (t-2-hex) promotes apoptotic signaling through coordinated effects on endoplasmic reticulum (ER) and mitochondrial function in *Saccharomyces cerevisiae*. We show that t-2-hex induces ER stress and activates the unfolded protein response via the canonical Ire1-Hac1 pathway, while simultaneously impairing mitochondrial protein import. Mechanistically, we identify the ER chaperone Kar2/BiP as a direct target of covalent lipidation by t-2-hex at a conserved cysteine residue, resulting in defective protein folding and proteostasis collapse. We further define the ER-associated short-chain dehydrogenase/reductase Fmp52 as a critical protective factor that, together with the mitochondrial aldehyde dehydrogenase Hfd1, counteracts lipid-induced stress. Notably, the human ortholog Tip30 functionally substitutes for Fmp52, highlighting evolutionary conservation. Our findings uncover a lipid-mediated mechanism that coordinately disrupts ER and mitochondrial proteostasis to drive apoptosis, and identify Fmp52/Tip30 as a conserved safeguard against lipid-induced cytotoxicity.

## Introduction

Mitochondria and the endoplasmic reticulum (ER) are central hubs of eukaryotic cell physiology, coordinating energy metabolism, protein trafficking, and quality control. Together, these organelles integrate signals that determine whether cells adapt to stress or undergo apoptosis. Mitochondria are well established as executors of cell death through release of cytochrome c and other pro-apoptotic factors ^1^, but the ER has emerged as a critical regulator of apoptosis by monitoring protein folding, calcium homeostasis, and lipid metabolism ^2–4^. Crosstalk between mitochondria and the ER is thus essential for maintaining proteostatic equilibrium and for controlling survival under stress ^5,6^.

Apoptosis plays a fundamental role in development and tissue homeostasis, and its deregulation contributes directly to human disease. Insufficient apoptosis underlies tumorigenesis, metastasis, and therapy resistance in cancer, while excessive or maladaptive apoptosis contributes to neurodegenerative diseases such as Alzheimer’s and Parkinson’s disease, as well as to inflammatory and metabolic disorders ^7–10^. Understanding the molecular mechanisms that control ER- and mitochondria-dependent apoptosis therefore has broad biomedical relevance.

Lipids are increasingly recognized as potent apoptotic regulators. Bioactive intermediates derived from sphingolipid catabolism are particularly important ^11,12^. The sphingolipid-derived aldehyde trans-2-hexadecenal (t-2-hex) is produced by sphingosine-1-phosphate lyase at the ER membrane and functions as a strong pro-apoptotic signal ^13–16^. The use of different model systems has recently identified direct targets of the lipid aldehyde in promoting apoptotic cell death. In mammalian cells, t-2-hex covalently modifies Bax to potentiate mitochondrial outer membrane permeabilization ^17,18^, whereas in yeast it targets the TOM import complex to inhibit mitochondrial protein import ^19^. Dedicated detoxifying enzymes such as the mitochondrial aldehyde dehydrogenase Hfd1 in yeast ^20^ are therefore required to counteract its toxicity ^15,19^. Importantly, in humans, defective detoxification of fatty aldehydes including t-2-hex due to mutations in the fatty aldehyde dehydrogenase (FALDH/ALDH3A2) causes the rare neurocutaneous disorder Sjögren-Larsson syndrome, characterized by ichthyosis, spasticity, and intellectual disability ^21,22^.

Chemoproteomic studies have revealed many potential protein targets of t-2-hex ^18,19^, suggesting broad cellular effects beyond mitochondria. The ER is of particular interest, as it is both the site of t-2-hex biogenesis and a central node of proteostasis. ER function relies on the Sec61/Sec63 translocation channel and the Kar2/BiP ATPase chaperone to ensure proper protein import and folding ^23^. These processes are tightly regulated by the unfolded protein response (UPR) and ER-associated degradation (ERAD) to prevent accumulation of misfolded proteins ^24^. Persistent ER stress can shift the UPR from adaptive to apoptotic signaling, linking ER quality control directly to cell death ^25^.

In this work, we use yeast as a model system to investigate how t-2-hex connects sphingolipid catabolism to ER dysfunction and apoptosis. We identify Fmp52, an ER-associated short-chain dehydrogenase/reductase (SDR) family member, as a novel protective factor against lipid-induced stress, and show that its human ortholog Tip30 functionally substitutes for this activity. Furthermore, we establish the essential ER chaperone Kar2/BiP as a direct lipidation target of t-2-hex. Our findings uncover a conserved mechanism by which bioactive lipids disrupt ER proteostasis and promote apoptosis, with implications for understanding the regulation of cell death in both normal physiology and disease.

## Results

### The pro-apoptotic lipid aldehyde t-2-hex causes ER-stress

Pro-apoptotic lipids such as the lipid aldehyde t-2-hex are generated within cells through the evolutionarily conserved sphingolipid degradation pathway (Figure 1A). Recent studies have revealed molecular details of t-2-hex activity at mitochondria, identifying Bax and components of the mitochondrial protein import machinery as direct lipid targets ^17,19^. In yeast, the dihydrosphingosine-1-phosphate lyase Dpl1, responsible for the final step in *t*-2-hex biogenesis, is an integral ER membrane protein with its catalytic domain oriented toward the cytoplasm ^26^. This localization suggests that t-2-hex is produced at the cytosolic face of the ER. In this work, we investigate the molecular targets and mechanisms that connect the pro-apoptotic function of t-2-hex with the ER. In our previous work, we used RNA-Seq to record the transcriptomic response of *hfd1* mutants, which accumulate elevated levels of *t*-2-hex ^19^. We specifically analyzed lipid-induced genes in this dataset for signatures of proteostatic ER stress, focusing on components of the ER-associated degradation (ERAD) pathway ^27^. Many ERAD-related gene functions were found to be significantly upregulated in response to *t*-2-hex (Figure 1B). To determine whether *hfd1* mutants exhibit phenotypes consistent with ER stress, we compared their growth to wild type under tunicamycin-induced ER stress conditions. Loss of Hfd1 function resulted in a significant hypersensitivity to tunicamycin (Figure 1C). Additionally, we determined that t-2-hex and tunicamycin inhibited cell proliferation in a synergistic manner, both in wild type and in *hfd1* mutant cells (Figure 1D). Next, we directly tested whether the pro-apoptotic lipid or the loss of the t-2-hex detoxifier Hfd1 induced ER stress, assessing activation of the unfolded protein response (UPR) using real-time luciferase reporters. As shown in Figure 1E, the UPRE-luciferase reporters revealed a clear dose-dependent activation in response to tunicamycin, which was strongly amplified in the absence of Hfd1. This was accompanied by both a higher inducibility (Figure 1F) and elevated steady-state activation of the UPR in *hfd1* mutants (Figure 1G). Direct treatment with t-2-hex elicited no detectable UPR reporter response in wild-type yeast, whereas *hfd1* mutants displayed a robust, dose-dependent activation (Figure 1H). Collectively, these findings demonstrate that t-2-hex induces ER stress, an effect further aggravated by the loss of Hfd1-mediated detoxification.

**Figure 1.**
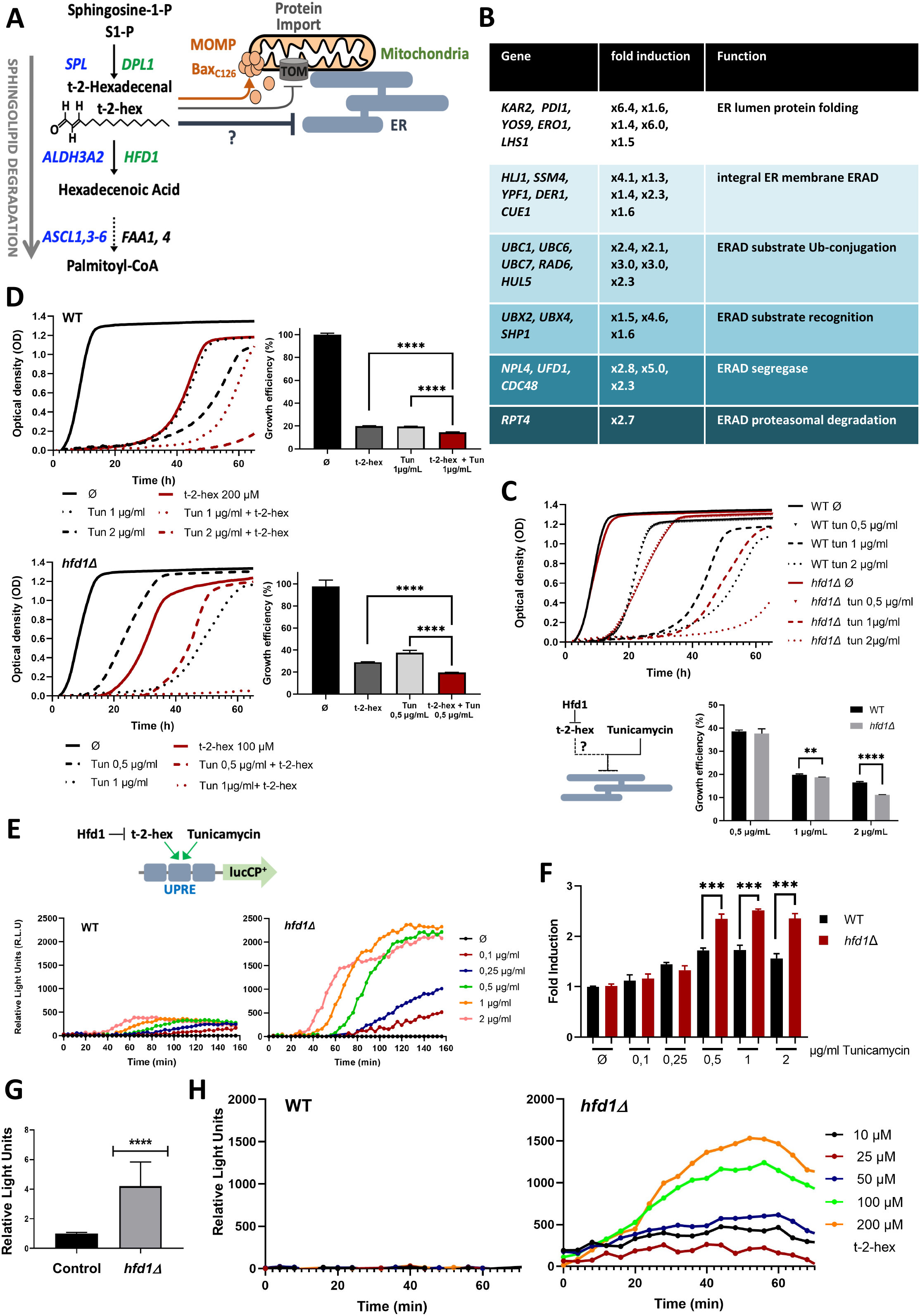
The pro-apoptotic lipid t-2-hex induces ER stress. (A) Overview of the generation of t-2-hex within the sphingolipid degradation pathway. Stress-activated enzymatic functions are depicted in green for yeast. The corresponding human enzymes involved in t-2-hex chemistry are shown in blue. t-2-hex disrupts mitochondrial function by various, recently discovered, mechanisms, such as C_126_ lipidation of human Bax or lipidation of the yeast TOM complex. (B) The ERAD pathway is induced by t-2-hex. The RNA Seq dataset of ^19^ was analyzed for ERAD pathway related gene functions. Significantly enriched ERAD genes are depicted together with their induction folds. (C) Loss of Hfd1 function leads to tunicamycin sensitivity. Quantitative growth assays of yeast wild type or *hfd1Δ* mutants in the presence of the indicated tunicamycin (tun) doses. Biological replicates n = 3. (D) t-2-hex and tunicamycin inhibit yeast growth synergistically. Quantitative growth assays of yeast wild type or *hfd1Δ* mutants in the presence of the indicated tunicamycin (tun) and/or t-2-hex (HD) doses. Biological replicates n = 3. (E) Tunicamycin induced gene expression via UPRE is enhanced by the loss of Hfd1 function. Quantitative live cell luciferase assays using a UPRE-luciferase reporter in yeast wild type or *hfd1Δ* mutants upon the indicated tunicamycin doses. Tunicamycin induction was corrected for mock treated cells for each concentration. Shown are mean values from 3 biological replicates. (F) Loss of Hfd1 function increases UPRE inducibility. Quantification of the fold induction by tunicamycin of UPRE comparing wild type and *hfd1Δ* mutants according to the results obtained in (E). (G) Basal activity of UPRE is enhanced by the loss of Hfd1 function. Steady state activity of the UPRE-luciferase reporter in the absence of tunicamycin stress comparing wild type (control) and *hfd1Δ* mutant. (H) t-2-hex induces UPRE mediated gene expression dependent on Hfd1 activity. Quantitative live cell luciferase assays using a UPRE-luciferase reporter in yeast wild type or *hfd1Δ* mutants upon the indicated t-2-hex doses. Shown are mean values from 3 biological replicates. ***p<0.001, ****p<0.0005 by Student’s unpaired t-test.

### t-2-hex induces the unfolded protein response via the Hac1-Ire1 pathway

In yeast, the unfolded protein response (UPR) to misfolded proteins in the ER lumen is mediated by the transmembrane sensor Ire1, which activates the transcription factor Hac1 through splicing of *HAC1* mRNA. This, in turn, initiates a transcriptional program that restores ER proteostasis ^28^. We next tested whether *t*-2-hex activates the UPR through this canonical pathway. We first measured intracellular Hac1 protein levels, which are typically transiently upregulated by known UPR activators such as tunicamycin or DTT (Figure 2A). Similarly, *t*-2-hex induced an initial increase in Hac1 protein abundance; however, this response appeared to be more sustained over time (Figure 2A). To further assess pathway activation, we quantified *HAC1* mRNA splicing in response to *t*-2-hex and confirmed that the lipid triggered Ire1-dependent splicing (Figure 2B). In contrast, treatment with the saturated analog *t*-2-hexadecanal (*t*-2-hex-H_2_), which retains the aldehyde group but lacks the ability to covalently modify protein targets ^18^, failed to induce *HAC1* mRNA splicing (Figure 2B). Together, these findings indicate that *t*-2-hex activates the UPR through the Ire1-Hac1 signaling pathway, most likely via direct modification of ER protein targets. We next asked whether UPR activation was required for tolerance to growth inhibition by t-2-hex. Comparing growth of yeast *ire1* and *hac1* mutants with wild type, we found that the Ire1/Hac1 UPR branch was dispensable for t-2-hex tolerance (Figure 2C). In a previous study, transposon mutagenesis identified genes potentially involved in tolerance to the pro-apoptotic lipid, including two ERAD components: the ER chaperone Eps1 and the ubiquitination factor Usa1 ^19^. Testing *eps1* mutants under t-2-hex stress revealed a significant sensitivity (Figure 2D), indicating that ERAD function is required for efficient tolerance, even though UPR activation itself is not.

**Figure 2.**
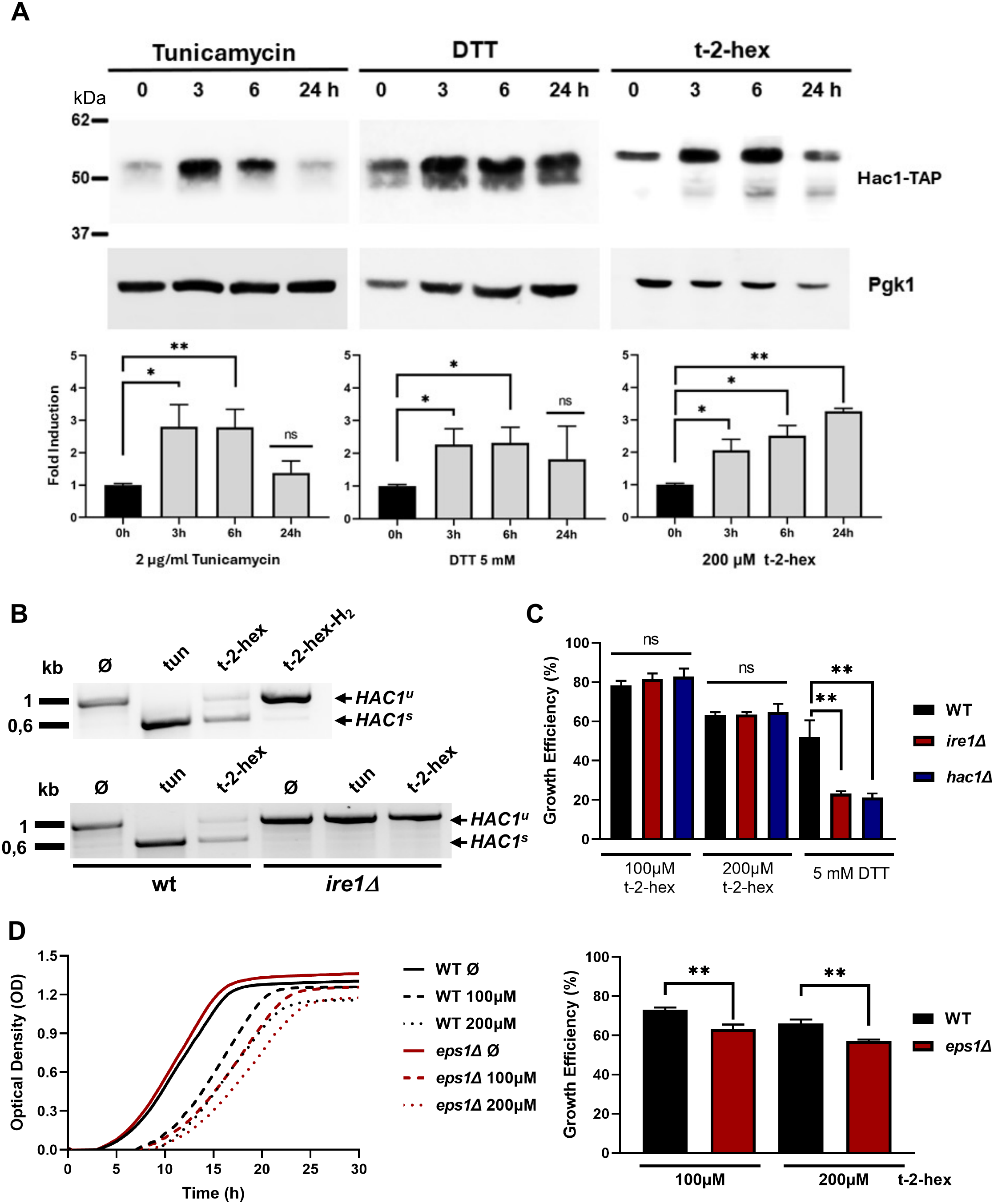
t-2-hex induces the unfolded protein response UPR via the canonical Hac1 pathway. (A) t-2-hex induces Hac1 protein levels. Hac1 was expressed as a Tap fusion from its chromosomal locus and cells were treated or not with the indicated doses of tunicamycin, DTT or t-2-hex for the indicated times. Hac1 protein abundance was quantified by anti-Tap western blot (upper panel) and quantified relative to uninduced levels (lower panel). Pgk1 protein levels served as loading control. (n = 3). (B) t-2-hex induces splicing of the *HAC1* mRNA. The appearance of unspliced (*HAC1^u^*) or spliced (*HAC1^s^*) mRNA was visualized by reverse transcriptase assays in yeast wild type or *ire1Δ* mutants. Cells were treated for 2h with tunicamycin (2μg/ml), t-2-hex (200μM) or the saturated analogue t-2-hex-H_2_ (200μM). (C) Activation of the UPR is not required for t-2-hex tolerance. Quantitative growth assays of yeast wild type, *ire1Δ* or *hac1Δ* mutants in the presence of the indicated DTT or t-2-hex doses. Biological replicates n = 3. (D) UPR function is important for t-2-hex tolerance. Quantitative growth assays of yeast wild type or *eps1Δ* mutants in the presence of the indicated t-2-hex doses. Biological replicates n = 3. **p<0.01 by Student’s unpaired t-test.

### Fmp52 functions in synergy with the Hfd1 dehydrogenase to promote t-2-hex detoxification

We previously performed transposon mutagenesis to identify, on a genomic scale, gene functions that promote tolerance to the pro-apoptotic lipid t-2-hex ^19^. Here, we reanalyzed these datasets focusing on ER-associated genes with potential roles in t-2-hex detoxification. Among them, *FMP52* showed the strongest under-enrichment signal, indicating that its loss impaired tolerance (Figure 3A). To test its role in detoxification, we evaluated the growth of *fmp52* deletion mutants in combination with loss of *HFD1*, the only known t-2-hex detoxifier, using quantitative growth assays (Figure 3B). Loss of *FMP52* function caused hypersensitivity to t-2-hex, comparable to that of *hfd1* mutants. Notably, the *fmp52 hfd1* double knockout exhibited markedly enhanced lipid sensitivity, revealing strong synergy between the two proteins in cellular resistance (Figure 3B). We then asked whether *FMP52* gene dosage influenced tolerance. To test this, we generated a constitutive overexpression strain via genomic promoter swapping and found that increased *FMP52* expression significantly improved tolerance (Figure 3C).

**Figure 3.**
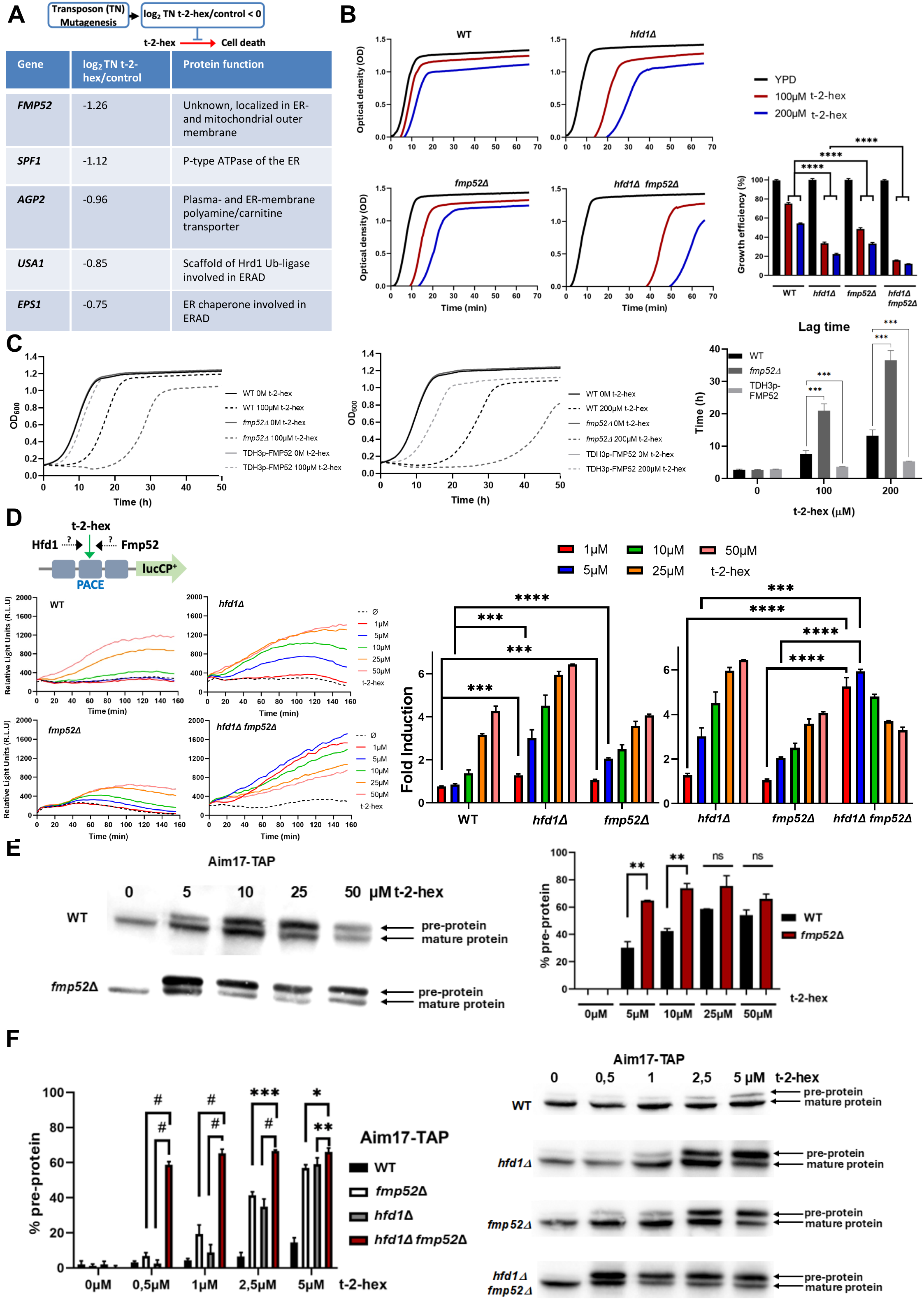
Fmp52 synergizes with Hfd1 in t-2 hex detoxification. (A) A saturated transposon screen SATAY was performed to identify underenriched gene functions with a potential role in t-2-hex tolerance and detoxification ^19^. Genes encoding ER proteins with a log_2_ ratio <-0.75 are depicted. (B) Fmp52 and Hfd1 are synergistically involved in t-2-hex tolerance. Quantitative growth assays of yeast strains harboring the indicated gene deletions in the presence of the indicated t-2-hex doses (left panels). Comparison of the respective growth efficiencies upon t-2-hex stress depending on *fmp52* and/or *hfd1* deletions (right panel). Biological replicates n = 3. (C) FMP52 gene dose determines t-2-hex tolerance. Left and middle panel: Quantitative growth assays of yeast wild type, *fmp52* deletion and constitutive *FMP52* overexpression (TDH3p-*FMP52*) in the presence of the indicated t-2-hex concentrations. Right panel: Comparison of the t-2-hex dependent lag phase. Biological replicates n = 3. (D) Hfd1 and Fmp52 synergistically control proteostasis signaling upon t-2-hex overload. Induction of a 3xPACE-lucCP^+^ reporter in wild type and indicated mutant yeast cells upon increasing t-2-hex doses. Left panels: Dose-response curves. Right panels: Maximal induction fold for each lipid dose tested (n = 3). (E) Loss of Fmp52 function increases t-2-hex induced mitochondrial pre-protein accumulation. Left panel: The appearance of unimported Aim17 mitochondrial precursor protein was induced by t-2-hex in chromosomally Tap-tagged wild type or *fmp52* mutant strains by anti-Tap western blot. Right panel: Aim17 pre-protein quantification. Biological replicates n = 3. (F) Fmp52 and Hfd1 have synergistic effects on mitochondrial pre-protein accumulation upon pro-apoptotic lipid stress. Right panel: t-2-hex induced pre-protein appearance in Aim17-Tap expressing strains of the indicated genetic background. Left panel: Aim17 pre-protein quantification. Biological replicates n = 3. *p<0.05, **p<0.01, ***p<0.001, ^#^p<0.0005 by Student’s unpaired t-test.

To determine whether the increased lipid sensitivity of the detoxification mutants was associated with enhanced cell death, we quantified membrane integrity by propidium iodide staining and apoptotic cell death by TUNEL assay. In wild-type cells, exposure to t-2-hex at concentrations up to 100 µM caused only minor cell death and no detectable TUNEL-positive cells. In contrast, loss of *HFD1* or *FMP52* progressively increased PI-positive cells upon lipid exposure, with the strongest effect observed when both detoxification pathways were disrupted (Supplementary Figure 1A). Importantly, t-2-hex also induced TUNEL-positive cells specifically in the detoxification mutants, reaching approximately 8% following treatment with 100 µM t-2-hex (Supplementary Figure 1B). These results demonstrate that Hfd1 and Fmp52 protect cells from t-2-hex-induced cell death and, importantly, prevent the execution of apoptotic cell death under conditions of lipid overload.

The dominant effect of t-2-hex in yeast is disruption of proteostasis, leading to strong PACE-dependent activation of proteasomal genes ^19^. To examine how Fmp52 and Hfd1 contribute to resistance, we assayed PACE-driven luciferase reporters in mutant strains. Loss of either factor moderately increased PACE activation, while the double mutant showed a dramatic, dose-dependent response, with minimal lipid exposure (1 µM) triggering maximal activation (Figure 3D). These results identify Fmp52 and Hfd1 as coordinated, essential detoxification factors that prevent proteostatic collapse under lipid stress. We next examined the recently described inhibition of mitochondrial pre-protein import by t-2-hex ^19^. Fmp52 prevented accumulation of unimported precursors at moderate lipid concentrations (Figure 3E), consistent with previous findings for Hfd1. Strikingly, the double mutant showed complete import inhibition at sub-µM lipid levels (Figure 3F), establishing Fmp52 and Hfd1 as essential surveillance factors for mitochondrial protein homeostasis under apoptotic lipid stress.

### Fmp52 is an ER-associated SDR protein controlling the unfolded protein response upon lipid stress

*FMP52* encodes a 231-amino acid protein structurally related to the short-chain dehydrogenase/reductase (SDR) family. SDR proteins are NAD(P)(H)-dependent oxidoreductases characterized by a Rossmann-fold scaffold that mediates dinucleotide binding ^29^. As shown in Figure 4A, Fmp52 harbors the glycine-rich consensus (GxxGxxG) within the cofactor-binding motif and a tyrosine-based catalytic center with an adjacent lysine (YxxxK), features typical of the “intermediate” SDR subfamily ^29^. AlphaFold structural predictions confirmed the presence of a Rossmann fold consisting of a central seven-stranded β-sheet flanked by four α-helices (Figure 4B). *FMP52* has a human ortholog, HTATIP2/TIP30, and sequence alignment revealed conservation of both the cofactor-binding and catalytic motifs (Figure 4A).

**Figure 4.**
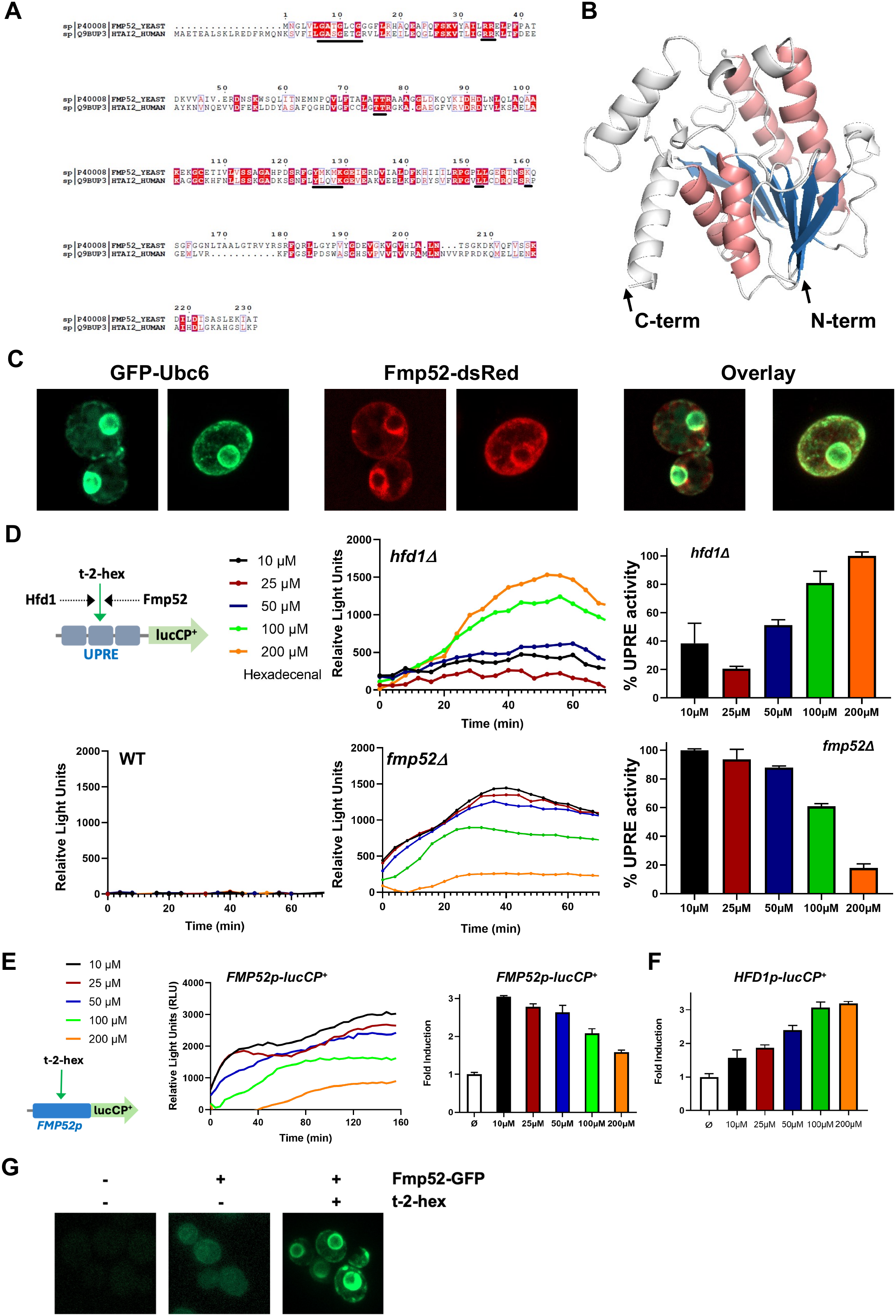
Fmp52 is an ER-associated protein that regulates the UPR in response to t-2 hex stress. (A) Fmp52 is a member of the short-chain dehydrogenase/reductase (SDR) family. Protein sequence comparison of Fmp52 with the human SDR protein HTATIP2/Tip30. Fmp52 (Uniprot 40008) and HTATIP2/TIP30 (Uniprot Q9BUP3) are depicted in the upper and lower line respectively. The core consensus sites in the NADP(H) binding motif (GxxGxxG) and in the active site (YxxxK) are highlighted, together with several other conserved motifs in the cofactor binding motif. (B) Fmp52 structure prediction by AlphaFold showing the characteristic distribution of central β-sheets (blue) combined with lateral α-helices (pink) of the Rossmann fold. Additional protein portions are depicted in grey. (C) Fmp52 localizes to the ER in yeast cells. Fmp52-dsRed and GFP-Ubc6 (ER marker) overexpressing yeast wild type cells were analyzed by confocal fluorescence microscopy. (D) Loss of Fmp52 function sensitizes UPR signaling upon t-2-hex stress. Quantitative live cell luciferase assays using a UPRE-luciferase reporter in yeast wild type or *hfd1Δ* and *fmp52Δ* mutants upon the indicated t-2-hex doses. Data were normalized for the mock treated control. Left panels: Continuous dose response curves. Right panels: Comparison of the maximal UPRE activity upon different lipid doses. Shown are mean values from 3 biological replicates. (E) The *FMP52* promoter is sensitively induced by t-2-hex. Quantitative live cell luciferase assays using a *FMP52*-luciferase reporter in yeast wild type cells. Dose responses were recorded continuously upon increasing t-2-hex concentrations (middle panel) and the maximal fold induction calculated for each lipid dose (right panel). Biological replicates n = 3. (E) Maximal fold induction of a *HFD1*-luciferase reporter in yeast wild type cells upon the same t-2-hex concentrations as in (D). Biological replicates n = 3. (G) Stress induced endogenous Fmp52 shows ER association. Strains expressing genomically GFP-tagged *FMP52* or the corresponding untagged control strain were analyzed by confocal fluorescence microscopy. Where indicated, cells were treated overnight with 100 μM t-2-hex prior to imaging.

To define the role of Fmp52 in apoptotic lipid detoxification, we first examined its intracellular localization. Endogenous GFP tagging did not yield clear results likely due to low protein expression, but overexpression of Fmp52-dsRed revealed a distinct ER localization, confirmed by colocalization with the ER marker GFP-Ubc6 (Figure 4C). We then assessed its impact on the unfolded protein response (UPR). UPR-specific luciferase assays revealed that loss of Fmp52 triggered strikingly stronger UPR activation at much lower t-2-hex concentrations (Figure 4D), highlighting a critical role for Fmp52 in ER protection that surpassed Hfd1. To further define the threshold for ER stress under conditions of impaired detoxification, we extended these reporter assays to the *hfd1Δ fmp52Δ* double mutant. Notably, t-2-hex concentrations as low as 0.5 μM elicited a robust UPR response (Supplementary Figure 2A), indicating that when both detoxification pathways are absent, even very low concentrations of exogenous t-2-hex are sufficient to compromise ER homeostasis. We next asked whether increased endogenous lipid production was sufficient to induce ER stress. To this end, we overexpressed *DPL1*, encoding the ER-localized sphingosine-1-phosphate lyase responsible for t-2-hex biogenesis, and monitored UPRE-luciferase activity. *DPL1* overexpression caused little UPR activation in wild type cells, whereas the response progressively increased upon loss of lipid detoxification capacity, with a modest increase in *hfd1Δ* cells and significant UPR activation in *hfd1Δ fmp52Δ* cells (Supplementary Figure 2B). These findings demonstrate that enhanced flux through the endogenous sphingolipid degradation pathway can compromise ER homeostasis when lipid detoxification is impaired, further supporting a physiological role for Fmp52 and Hfd1 in protecting the ER from endogenous lipid stress.

We next asked whether *FMP52* itself is regulated by lipid stress. A luciferase reporter under control of the *FMP52* promoter showed strong induction by t-2-hex, with high sensitivity to low micromolar concentrations (Figure 4E). In direct comparison, the *HFD1* reporter required higher lipid levels for activation, underscoring Fmp52 as the more responsive regulator (Figure 4F).

Having established that *FMP52* expression is induced by t-2-hex, we next examined the intracellular localization of endogenous Fmp52 under lipid stress conditions. Using a strain expressing genomically GFP-tagged Fmp52, we observed a weak and diffuse fluorescence signal under basal conditions, with only limited overlap with the ER. Following prolonged exposure to t-2-hex, Fmp52 adopted a pronounced ER-like localization pattern, consistent with enhanced association of endogenous Fmp52 with the ER (Figure 4G).

### Fmp52 is required for ER protein maturation under t-2-hex stress

To investigate the role of Fmp52 in ER homeostasis during lipid stress, we monitored client protein import using a Gal-inducible Pdi1-DHFR “clogger” fusion protein ^30^. This construct generates a translocation challenge due to the stable DHFR fold and enables direct visualization of ER import and glycosylation states by western blot.

In wild type cells, steady state expression of the clogger resulted in complete import and glycosylation, unaffected by the presence of t-2-hex. By contrast, *ste24Δ* cells accumulated unglycosylated and hemiglycosylated forms, consistent with defective translocon function; tunicamycin treatment completely abolished glycosylation (Figure 5A). We next compared wild type and *fmp52Δ* strains under rapid clogger induction. While wild type cells efficiently processed the protein, *fmp52Δ* cells showed incomplete maturation, suggesting a requirement for Fmp52 in maintaining ER import capacity (Figure 5B). To test this more stringently, we analyzed clogger maturation during the early hours of induction under lipid stress. Strikingly, Pdi1-DHFR processing was selectively impaired by t-2-hex in *fmp52Δ* cells but remained unaffected by the saturated analogue t-2-hex-H_2_ (Figure 5C). To exclude a general effect of t-2-hex on Gal-induced expression, we monitored *GAL1p*-driven luciferase activity during early induction in live cell assays. Luciferase expression remained unaffected by the lipid, regardless of Fmp52 presence (Figure 5C). We alternatively examined the endogenous ER client Gas1, a GPI-anchored protein requiring ER processing for cell wall delivery. Gas1 maturation was transiently blocked by t-2-hex, specifically by its unsaturated form (Figure 5D). These findings indicate that Fmp52 is essential to maintain ER client maturation under unsaturated lipid stress.

**Figure 5.**
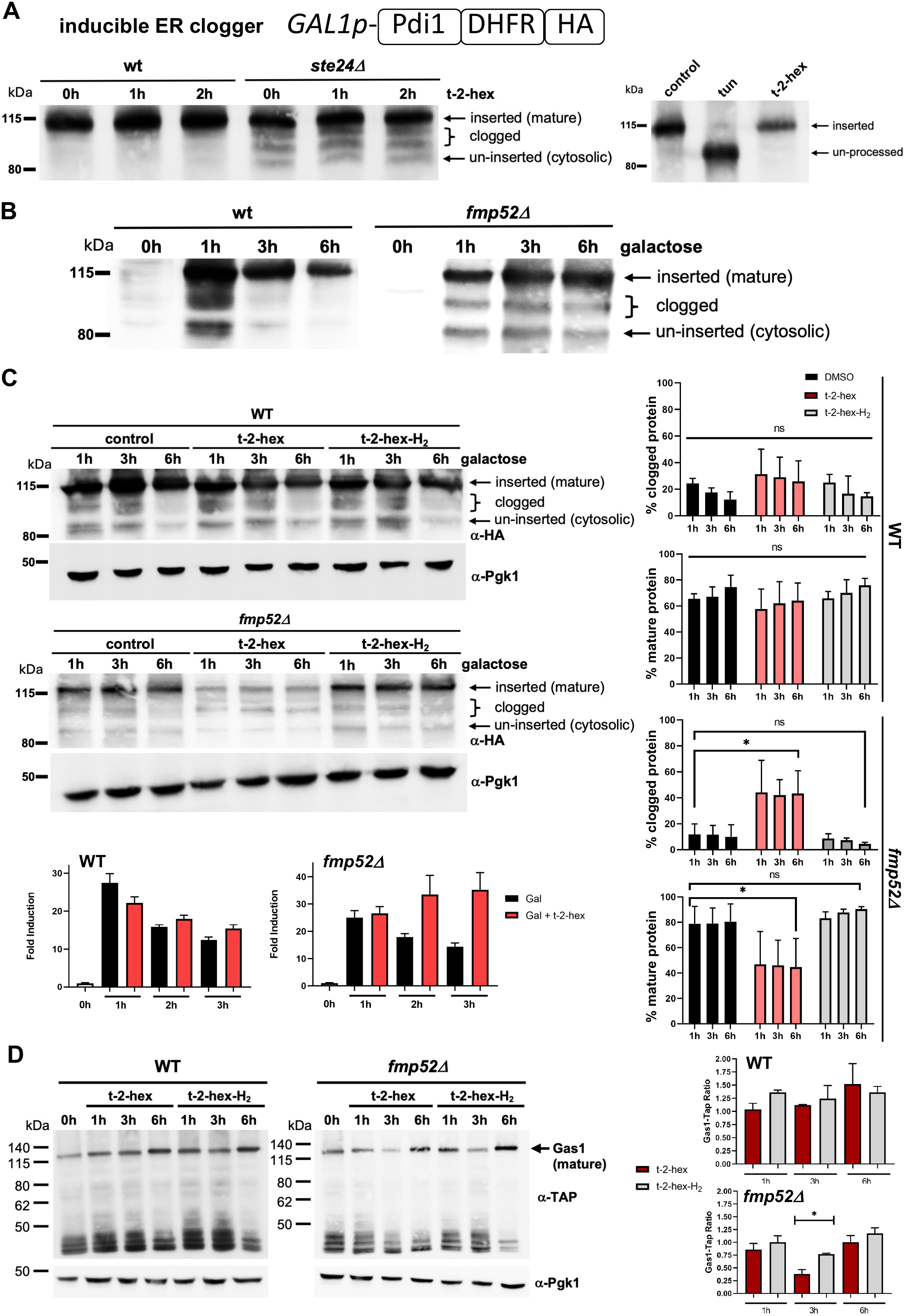
Fmp52 controls maturation of ER proteins upon pro-apoptotic lipid stress (A) An inducible synthetic clogger protein (Pdi1-DHFR) was used to monitor protein processing at the ER. The Pdi1-DHFR clogger was induced by galactose over night in yeast wt and *ste24Δ* mutants and mature and different un-processed forms visualized by anti-HA blotting upon the indicated t-2-hex treatment (left panel). Tunicamycin (2μg/ml) and t-2-hex (200μM) treatment over night of Pdi1-DHFR clogger expressing wild type cells (right panel). (B) Induced expression of the Pdi1-DHFR clogger in yeast wt and *fmp52Δ* mutants. (C) Fmp52 is necessary for Pdi1-DHFR clogger processing upon t-2-hex stress. Upper left panels: The Pdi1-DHFR clogger was induced for the indicated times in yeast wt and *fmp52Δ* mutants in the presence or not of 200μM of t-2-hex or t-2-hex-H_2_. Pgk1 was used as a loading control. Right panels: Quantification of the percentage of mature and clogged Pdi1-DHFR protein relative to total protein. Biological replicates n = 3. Lower left panels: Cytosolic luciferase was galactose induced in yeast wt and *fmp52Δ* mutants in the presence or not of 200μM of t-2-hex and expression monitored by live cell luciferase assays. Biological replicates n = 3. (D) Fmp52 is necessary for endogenous Gas1 maturation upon t-2-hex stress. Left panels: Gas1-TAP fusion proteins were endogenously expressed in yeast wt and *fmp52Δ* mutants and protein expression and maturation visualized by anti-TAP blotting. Pgk1 was used as a loading control. Cells were treated with 200μM of t-2-hex or t-2-hex-H_2_. Right panels: Quantification of the ratio of Gas1 mature protein relative to total protein (Pgk1). Biological replicates n = 3. *p<0.05, ns = not significant by Student’s unpaired t-test.

### Kar2/BiP is a direct lipidation target of t-2-hex, and its C_63_ residue contributes to lipid-stress tolerance

Given the role of Fmp52 in counteracting lipid-induced ER stress, we next asked what underlies t-2-hex-triggered dysfunction. T-2-hex is known to covalently modify proteins preferentially at specific cysteine residues ^18^, such as Bax in human mitochondria and TOM subunits in yeast ^17,19^. To explore potential ER-specific targets, we re-analyzed our recent in vitro chemoproteomic dataset, which revealed several ER-associated proteins displaying different degrees of enrichment upon t-2-hex-alkyne labeling (Figure 6A). Among these, we selected Kar2, Spf1, Sec61 and Sec63 for targeted validation. While Sec61 and Sec63 reached statistical significance in the original chemoproteomic dataset, Kar2 and Spf1 showed enrichment without passing the significance threshold. Spf1 is a non-essential ER ion transporter involved in Ca²⁺ homeostasis and general ER function, whereas Kar2 (BiP in mammals) is an essential ATPase chaperone that facilitates protein import and folding in the ER through its interaction with the Sec62/Sec63 translocation complex. Targeted lipidation assays in yeast extracts revealed robust enrichment of Kar2 at low micromolar concentrations of t-2-hex (25 μM) (Figure 6B). Under the same assay conditions, weak enrichment was observed for Spf1, whereas no detectable enrichment was obtained for Sec61 or Sec63 (Figure 6C). Because detection in this targeted assay may be influenced by the abundance and recovery of the individual TAP-tagged proteins, the absence of detectable Sec61 or Sec63 enrichment does not exclude their modification by t-2-hex. To determine whether Kar2 is modified by t-2-hex in living cells, we next performed the lipidation assay directly in intact yeast. Cells were exposed to t-2-hex-alkyne prior to lysis, extensively washed to remove extracellular probe, and subjected to click chemistry and streptavidin enrichment. Specific Kar2 lipidation was detected across a range of low t-2-hex concentrations (10-50 μM) (Figure 6D). These results demonstrate that Kar2 is covalently modified by t-2-hex in intact yeast cells and support its physiological relevance as a target of the lipid aldehyde.

**Figure 6.**
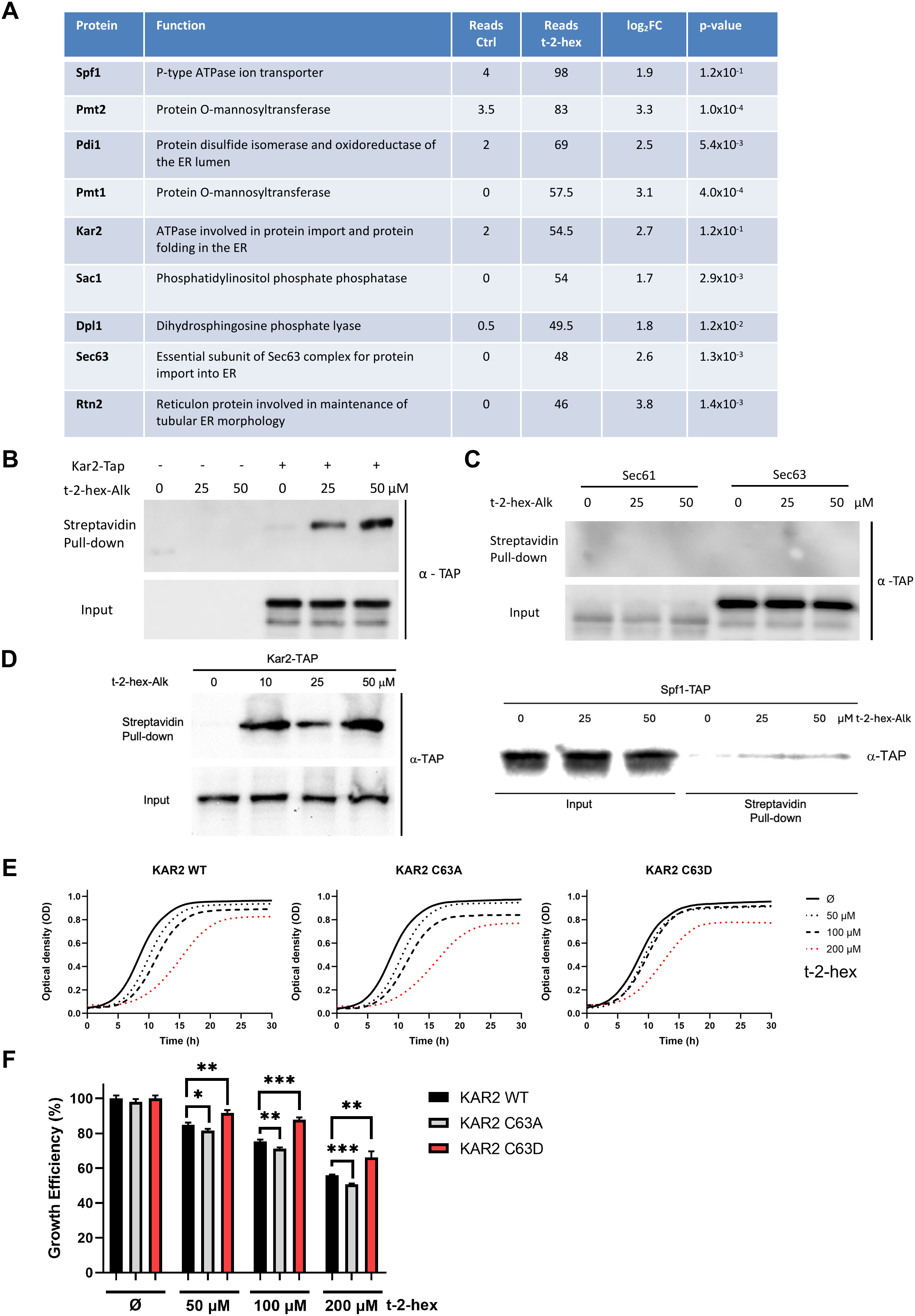
Kar2 is a direct and physiologically relevant lipidation target for t-2-hex. (A) Re-analysis of the previously published t-2-hex-alkyne chemoproteomic dataset ^19^ for ER-associated candidate proteins. The table shows mean spectral reads in mock-treated and t-2-hex-alkyne-treated samples (n = 2), together with log2 fold enrichment and adjusted p-values. Proteins are ranked according to spectral abundance in the lipid-treated samples. Statistical significance in the original discovery dataset is indicated by the corresponding adjusted p-values. (B) Kar2 is lipidated *in vitro* by t-2-hex. Upper panel: t-2-hex-Alkyne was used in the indicated concentrations to lipidate proteins in enriched ER/mitochondrial preparations from Kar2-Tap expressing cells or control cells. After t-2-hex addition, input samples were generated directly, while pull-down samples were treated with click chemistry for covalent biotin linkage and subsequent Streptavidin purification. Kar2 was detected in all samples by anti-Tap western blot. (C) Analysis of t-2-hex in vitro lipidation of the ER translocon subunits Sec61 and Sec63 and the Spf1 ATPase. Lipidation assays were performed as in (B) using the indicated Tap fusion strains. (D) Kar2 is lipidated in vivo by t-2-hex. t-2-hex-Alkyne was used in the indicated concentrations on intact cells. Input samples were generated directly after cell lysis, while pull-down samples were treated with click chemistry for covalent biotin linkage and subsequent Streptavidin purification. (E) Kar2 C_63_ is important for t-2-hex tolerance. Yeast strains expressing wild type or point mutated (C_63_A, C_63_D) versions of Kar2 were analyzed for t-2-hex tolerance by quantitative growth assays in the presence of the indicated lipid doses. (E) Comparison of the growth efficiency upon t-2-hex stress dependent on Kar2 C_63_. Biological replicates = 3. *p<0.05, **p<0.01, ***p<0.001 by Student’s unpaired t-test.

Kar2 contains a unique cysteine at position 63 (C63), previously shown to be chemically modified and essential for ER protein folding under oxidative stress ^31,32^. Oxidation of C63 converts Kar2 into a more active chaperone, although this residue is dispensable for normal growth ^31^. To assess the functional contribution of Kar2 C63, we used previously established Kar2 replacement strains, genetically different from the BY4741 background used for the other experiments in this study, in which the endogenous *KAR2* gene is deleted and viability is maintained by plasmid-borne WT *KAR2* or the C63A or C63D variants: C63A, which cannot switch to the active form, and C63D, which mimics the oxidized, active chaperone ^31^. Both mutants, lacking cysteine lipidation, were tested for tolerance to t-2-hex (Figure 6E). Alanine substitution of Kar2-C63 reduced lipid tolerance, whereas the Aspartic acid substitution markedly enhanced it (Figure 6F). These results indicate that the functional state of Kar2 C63 influences the cellular response to t-2-hex stress. Together with the direct detection of Kar2 lipidation in intact cells, they support a model in which covalent modification of Kar2 contributes to t-2-hex-induced ER dysfunction. However, these experiments do not by themselves establish C63 as the site of t-2-hex modification.

### Human Tip30 is a functional homolog of yeast Fmp52 in t-2-hex tolerance

Tip30, the human ortholog of yeast Fmp52, is a tumor suppressor that restrains proliferation, inhibits metastasis, and promotes apoptotic cell death ^33,34^. Although no specific metabolic substrate has been identified to date, Tip30 adopts the typical fold of short-chain dehydrogenase/reductase (SDR) proteins ^35^. To assess its role in lipid detoxification, we expressed Tip30-Flag in wild-type and t-2-hex-sensitive yeast strains (Figure 7A). Tip30 fully complemented the loss of Fmp52, restoring wild-type growth across all tested t-2-hex concentrations (Figures 7B,C). Complementation also suppressed t-2-hex-induced UPR activity in *fmp52Δ* cells, as Tip30 expression strongly reduced UPRE-luciferase activation (Figure 7D). Moreover, Tip30 alleviated lipid-mediated mitochondrial defects, markedly reversing Aim17 precursor accumulation in *hfd1Δfmp52Δ* cells (Figure 7E). Finally, under general apoptotic stress, Tip30 expression significantly improved yeast resistance to acetic acid-induced growth inhibition (Figure 7F).

**Figure 7.**
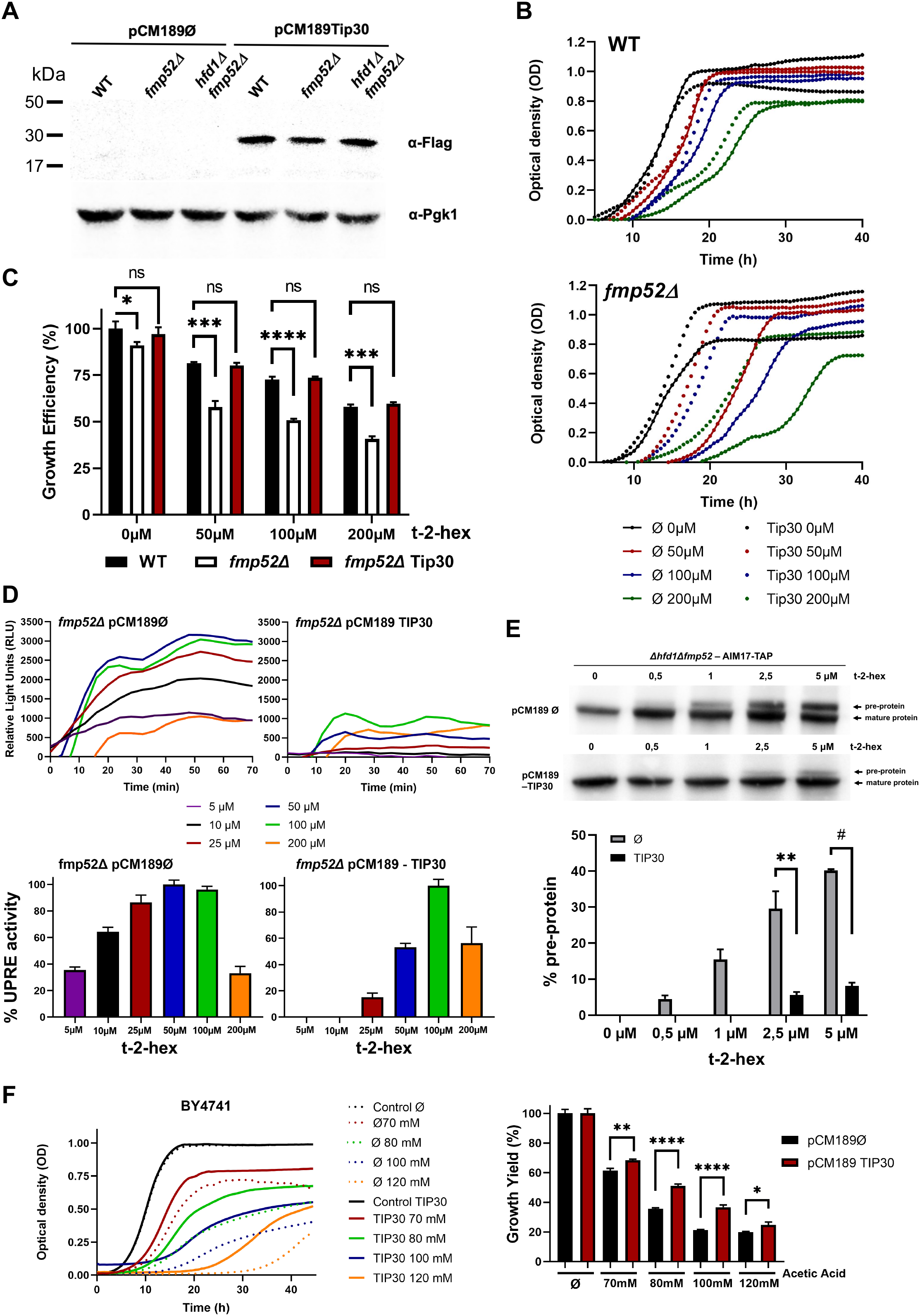
Human Tip30 is a functional homolog of yeast Fmp52 in t-2-hex tolerance. (A) Human Tip30-Flag was constitutively expressed in yeast wild type and the indicated *fmp52* mutants. Tip30 was detected in yeast total extracts by anti-Flag blotting. Pgk1 was used as a loading control. (B) Tip30 complements t-2-hex induced growth inhibition. Quantitative growth assays of yeast wild type or *fmp52Δ* mutants expressing human Tip30 or not in the presence of the indicated t-2-hex doses. Biological replicates n = 3. (C) Comparison of the t-2-hex dependent inhibition of the continuous growth experiments in (B). Biological replicates n = 3. (D) Tip30 expression reduces UPR activation upon t-2-hex stress. Quantitative live cell luciferase assays using a UPRE-luciferase reporter in yeast *fmp52Δ* mutants, complemented or not by Tip30 expression, upon the indicated t-2-hex doses. Upper panel: Continuous dose response curves. Lower panel: Comparison of the t-2-hex dependent UPRE activation. Shown are mean values from 3 biological replicates. (E) Tip30 activity reduces mitochondrial protein import defects upon t-2-hex overload. Upper panel: The appearance of unimported Aim17 mitochondrial precursor protein was induced by t-2-hex in chromosomally Tap-tagged *Δfmp52Δhfd1* mutant strains and visualized by anti-Tap western blot. Tip30 was co-expressed or not as indicated. Lower panel: Aim17 pre-protein quantification relative to total Aim17. Biological replicates n = 3. (F) Tip30 protects yeast cells from pro-apoptotic stress caused by acetic acid. Yeast wild type cells expressing or not human Tip30 were treated with the indicated acetic acid concentrations. Left panel: Quantitative growth assays of three independent biological replicates. Right panel: Comparison of the Tip30 dependent growth efficiency. Biological replicates n = 3. *p<0.05, **p<0.01, ***p<0.001, ****p<0.0005 by Student’s unpaired t-test.

The ability of human Tip30 to functionally replace Fmp52 raised the possibility that the conserved SDR catalytic center contributes to t-2-hex detoxification. To address this question, we generated a Tip30 variant in which both residues of the conserved YxxxK catalytic motif were mutated. The resulting Tip30 catalytic-site mutant was expressed in the *hfd1Δ fmp52Δ* background and compared with wild-type Tip30 (Supplemental Figure 3A).

Whereas wild-type Tip30 efficiently restored resistance to t-2-hex, the catalytic-site mutant displayed a significantly reduced ability to rescue t-2-hex-induced growth inhibition (Supplementary Figure 3B and C). Importantly, the mutant retained partial complementation, indicating that disruption of the YxxxK motif does not completely abolish Tip30-dependent protection. These findings demonstrate that the conserved catalytic center of Tip30 contributes substantially to its protective function against t-2-hex, while also suggesting that catalytic-site-independent activities may contribute to the residual rescue.

Together, these results demonstrate that human Tip30 functionally substitutes for yeast Fmp52 and protects cells against the deleterious effects of t-2-hex. The reduced complementation observed upon disruption of the conserved YxxxK motif further indicates that the Tip30 catalytic center contributes to this protective activity.

## Discussion

Here we provide several lines of evidence supporting a novel model that integrates the molecular steps leading to ER dysfunction upon apoptotic lipid stress with the cellular mechanisms that prevent cell death under these conditions (Figure 8). Central to this apoptotic regulatory circuit is the bioactive lipid aldehyde t-2-hex, an intermediate of the stress-induced sphingolipid degradation pathway operating at the ER. Until now, our understanding of physiologically important targets of t-2-hex has been largely restricted to mitochondrial proteins, such as Bax and components of the TOM complex, where cysteine-specific lipidation interferes with mitochondrial outer membrane permeability or preprotein import ^17,19^. In this study, we focus on the inhibitory functions of the apoptotic lipid at the ER and identify the Kar2/BiP chaperone as a physiologically important lipidation target.

**Figure 8.**
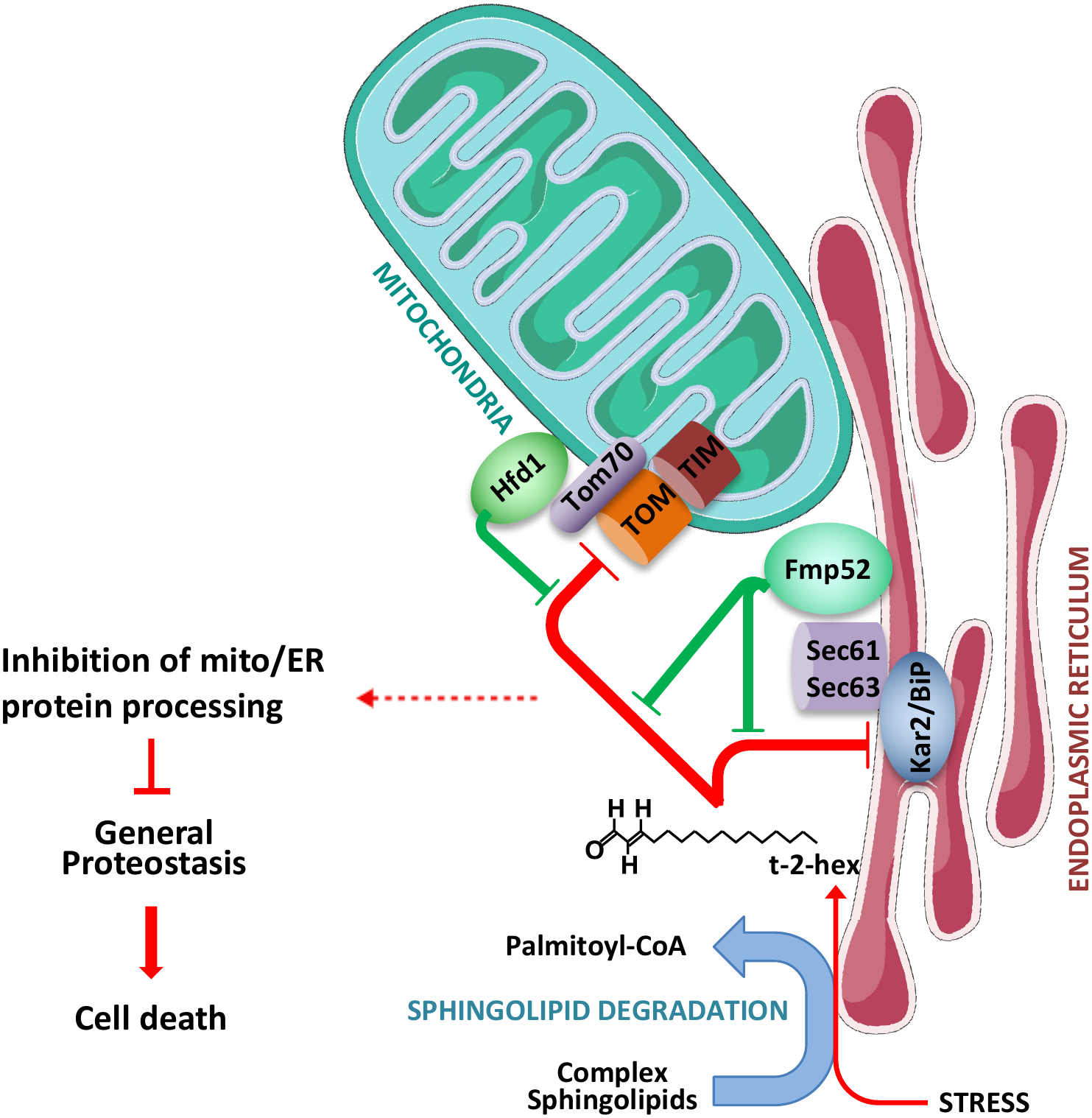
Model of the pro-apoptotic function of t-2-hex at the ER and mitochondria, and the anti-apoptotic roles of Fmp52 and Hfd1. Under stress conditions, the bioactive lipid aldehyde t-2-hex accumulates at the ER and targets essential protein functions, including the ER chaperone Kar2/BiP and the mitochondrial TOM complex. Lipidation of these factors disrupts ER and mitochondrial protein processing, leading to precursor accumulation and proteostatic imbalance that can trigger apoptosis if not properly resolved. The stress-inducible safeguard proteins Fmp52 and Hfd1 act synergistically from their respective organelles to detoxify t-2-hex and counteract its inhibitory effects. Pro-apoptotic pathways are indicated in red, and anti-apoptotic detoxification mechanisms in green.

Yeast Kar2 contains a single, evolutionarily conserved cysteine residue at position 63 within the ATP-binding domain of the chaperone. C63 plays a redox-active regulatory role in Kar2, and its site-specific oxidation during oxidative stress has been shown to enhance folding capacity at the ER ^31^. Large non-polar substitutions at C63 disrupt the essential chaperone function of Kar2 ^36^, suggesting that lipidation at this site could significantly impair Kar2/BiP basal activity and thereby compromise ER protein folding capacity. In the present study, we demonstrate that Kar2 is covalently lipidated by t-2-hex not only in cell extracts but also following treatment of intact yeast cells. This establishes that Kar2 is a bona fide cellular target of t-2-hex modification. The C63A and C63D substitution experiments further show that the regulatory state of this residue influences cellular tolerance to t-2-hex stress. However, because C63 is also a known redox-regulatory residue, the opposing phenotypes of C63A and C63D cannot by themselves distinguish between direct lipidation-dependent effects and effects arising from altered redox regulation of Kar2. We therefore interpret these experiments as evidence that Kar2 C63 contributes to the cellular response to t-2-hex, rather than as definitive evidence that C63 is the site of lipidation. Direct identification of the t-2-hex-modified residue(s) by mass spectrometry will be required to establish the precise site of modification.

Kar2/BiP is a central regulator of ER protein quality control, and its expression is strongly upregulated in response to apoptotic stimuli from yeast to humans ^37,38^. Its activity and availability critically shape the severity of ER stress ^39^. Consistent with this role, BiP has been shown to influence the execution of pro-apoptotic pathways: in mammalian systems, excessive ER stress induces mitochondria mediated cell death ^40^, BiP ablation in mouse models triggers neuronal apoptosis ^41^, repression of BiP enhances ER-mediated apoptosis in human cells ^42^, and elevated BiP levels can suppress apoptotic signaling ^43^. Our finding that Kar2/BiP is directly targeted by the pro-apoptotic lipid t-2-hex uncovers an additional regulatory layer acting on this key chaperone, providing new insight into how apoptotic stimuli modulate BiP function at the molecular level. Despite this strong connection, Kar2/BiP, although physiologically critical, is unlikely to be the sole ER target of t-2-hex, as demonstrated by the low levels of lipidation detected for the ER ATPase Spf1.

An important advance emerging from our work is the identification of Fmp52 as a previously unrecognized regulator of lipid-induced apoptosis. Fmp52 has not previously been associated with ER quality control, apoptotic signaling, or lipid detoxification, yet several lines of evidence now position it as a central node in the cellular response to t-2-hex. First, the protein localizes primarily to the ER, placing it at the site of lipid generation and ER proteostasis control. Second, loss of Fmp52 confers marked hypersensitivity to t-2-hex. Third, *fmp52Δ* cells exhibit exaggerated UPR activation under lipid stress and fail to maintain processing of both endogenous and synthetic ER client proteins. Finally, Fmp52 is essential to preserve mitochondrial protein import during t-2-hex exposure. Together, these observations reveal Fmp52 as a determinant of organellar proteostasis during apoptotic lipid stress. The protein acts as a protective factor that shields both ER and mitochondria from the proteotoxic consequences of t-2-hex accumulation. Importantly, loss of Fmp52 also increased both overall and TUNEL-positive apoptotic cell death upon t-2-hex exposure, providing direct evidence that its protective function extends beyond maintenance of growth and proteostasis to suppression of lipid-induced apoptosis. A central question raised by our findings concerns the precise molecular activity of Fmp52. *FMP52* is transcriptionally induced by t-2-hex and responds to lipid exposure with even higher sensitivity than *HFD1*, the previously established mitochondrial t-2-hex dehydrogenase ^15,20^. Its predicted enzymatic architecture, possessing both the classical Rossmann-fold and the conserved catalytic motifs of the SDR superfamily, raises the possibility that Fmp52 is itself a lipid-modifying enzyme. Its lack of predicted transmembrane segments suggests that Fmp52 associates peripherally with the ER membrane, where it could act directly on t-2-hex or its reactive intermediates. However, whether Fmp52 directly metabolizes the lipid remains unresolved. This point is particularly intriguing given that its human ortholog Tip30 has been proposed to act primarily as a regulatory rather than enzymatic SDR protein, largely due to atypical dimerization behavior and the lack of substrate candidates ^29,35^. Our demonstration that human Tip30 rescues all major phenotypes of *fmp52Δ* cells, including UPR hyperactivation, mitochondrial import defects, and t-2-hex hypersensitivity, confirms the evolutionary conservation of this protective mechanism, but does not yet distinguish whether either protein is a bona fide lipid dehydrogenase. The mutational analysis of human Tip30 provides additional evidence that the conserved SDR catalytic center contributes to this protective function. Disruption of both residues within the conserved YxxxK motif reduced the ability of Tip30 to complement the t-2-hex sensitivity of *hfd1Δ fmp52Δ* cells. Thus, the protective activity of Tip30 is at least partly dependent on its conserved catalytic center. However, the residual rescue observed with the catalytic-site mutant indicates that catalytic-site-independent functions may also contribute to Tip30-mediated protection. Moreover, because the present study does not directly demonstrate enzymatic conversion of t-2-hex by purified Fmp52 or Tip30, these findings do not by themselves establish that either protein directly metabolizes t-2-hex. Biochemical characterization will be required to determine whether the SDR domain catalyzes t-2-hex reduction or oxidation and to define its substrate specificity.

Integrating our current findings with earlier observations allows us to expand the mechanistic framework of lipid-mediated apoptosis. We previously established that t-2-hex inhibits mitochondrial protein import through direct lipidation of TOM complex components ^19^. The present study now demonstrates that t-2-hex simultaneously impairs ER protein maturation by lipidating Kar2/BiP. Thus, the lipid acts through a dual-organelle inhibitory mechanism, compromising proteostasis at both major sites of protein translocation and folding. Importantly, loss of Fmp52 dramatically enhances this vulnerability: sub-micromolar concentrations of t-2-hex are sufficient to elicit complete mitochondrial import arrest when both Fmp52 and Hfd1 are absent. Importantly, the protective function of Fmp52 and Hfd1 is not restricted to experimentally added lipid. Increasing endogenous lipid production through *DPL1* overexpression was sufficient to activate the UPR when both detoxification pathways were compromised, indicating that endogenous sphingolipid catabolism can generate ER stress and that Fmp52 and Hfd1 normally buffer the cell against this intrinsic lipid burden. These synergistic perturbations indicate that Fmp52 and Hfd1 represent the two principal anti-apoptotic systems that buffer the cell against bioactive lipid accumulation.

The combined targeting of ER and mitochondria by t-2-hex is mechanistically significant for several reasons. First, both organelles act as proteostasis hubs whose coordination is essential for cellular viability ^44,45^. Inhibition at either site produces accumulation of unprocessed precursors in the cytosol, but concurrent inhibition amplifies the collapse of protein homeostasis. Second, ER stress and mitochondrial import stress are intimately connected. Specifically, correct targeting and maturation of mitochondrial preproteins requires ER function and the physical interconnection of both organelles ^46,47^. Moreover, defects in mitochondrial import are known to activate ER stress pathways, and mislocalized mitochondrial precursors are buffered at the ER ^48^. Conversely, ER proteins which fail to fold and mature correctly cause mitochondrial dysfunction ^49^. Thus, simultaneous targeting of Kar2/BiP and the TOM complex by t-2-hex establishes a self-reinforcing cycle of proteostatic failure. Third, collapse of ER-mitochondria proteostasis is increasingly recognized as a potent trigger of apoptosis and has been implicated in neurodegenerative disease and metabolic pathology ^50^. Our findings therefore support a model in which t-2-hex leverages organelle crosstalk to intensify apoptotic signaling.

Collectively, our data define a refined, integrated model of lipid-mediated apoptosis in which t-2-hex acts as a bifunctional inhibitor of ER and mitochondrial proteostasis, while Fmp52 and Hfd1 constitute spatially distributed, synergistic detoxification systems that counteract its cytotoxicity. By lipidating Kar2/BiP at the ER and targeting the TOM complex at mitochondria, t-2-hex induces proteostatic imbalance that drives the cell toward apoptosis unless neutralized by these protective factors. The capacity of human Tip30 to substitute for yeast Fmp52 further suggests that this regulatory circuit is conserved across eukaryotes and may be relevant to human pathologies associated with aldehyde detoxification defects. These insights lay the foundation for future studies aimed at dissecting the enzymatic potential of Fmp52/Tip30 and at exploring how lipid-driven proteostasis collapse contributes to disease.

## Materials and methods

### Yeast strains and growth conditions

The yeast strains used in this work are detailed in Table 1. Yeast cultures were grown, unless otherwise indicated, at 28°C in Yeast Extract Peptone Dextrose (YPD) media containing 2% glucose, Yeast Extract Peptone Galactose (YPGal) media containing 2% galactose or in Synthetic Dextrose (SD) media containing 0.67% yeast nitrogen base with ammonium sulphate and without amino acids, 50mM succinic acid (pH5.5) and 2% of the respective sugar. According to specific auxotrophies of individual strains, methionine (10mg/l), histidine (10mg/l), leucine (10mg/l) or uracil (25mg/l) were supplemented. Yeast cells were transformed with the lithium acetate/PEG method described by ^51^. Strains expressing particular proteins as Tap-fusions from the chromosomal locus are described in ^52^. Genomic deletion of particular genes was done with PCR based methods using KAN-MX, hph-MX or his5^+^ (*S. pombe*) containing plasmids pUG6, pAG32 or pUG27 ^53,54^. All deletion strains were verified with diagnostic PCR on purified genomic DNA. For genomic promoter swapping, a KAN-MX::TDH3p containing construct was made by cloning the *TDH3* promoter in the pUG6 plasmid at the *Eco*RV and *Spe*I sites. The Fmp52 constitutive overexpressor strain was made by replacing the native promoter with the KAN-MX::TDH3p cassette.

**Table 1.**
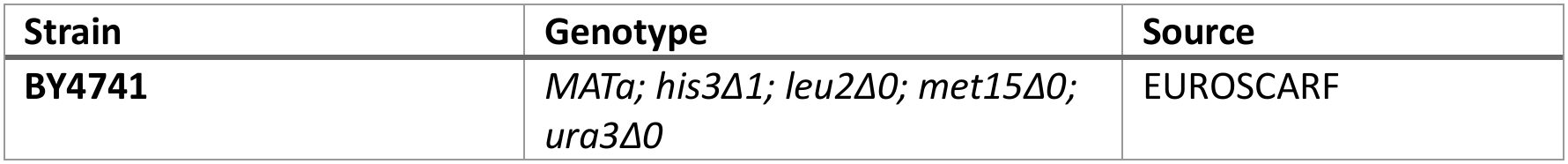
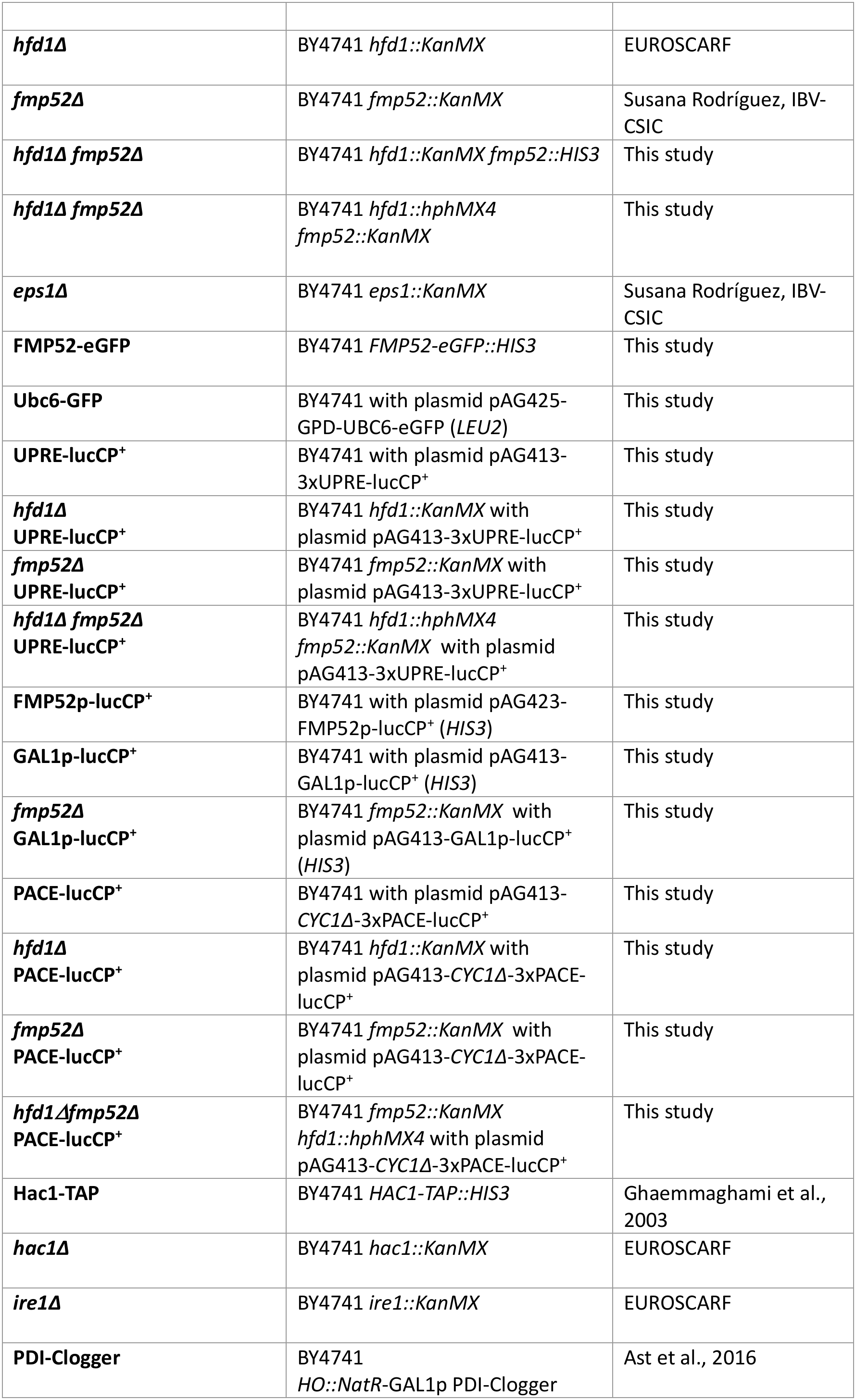
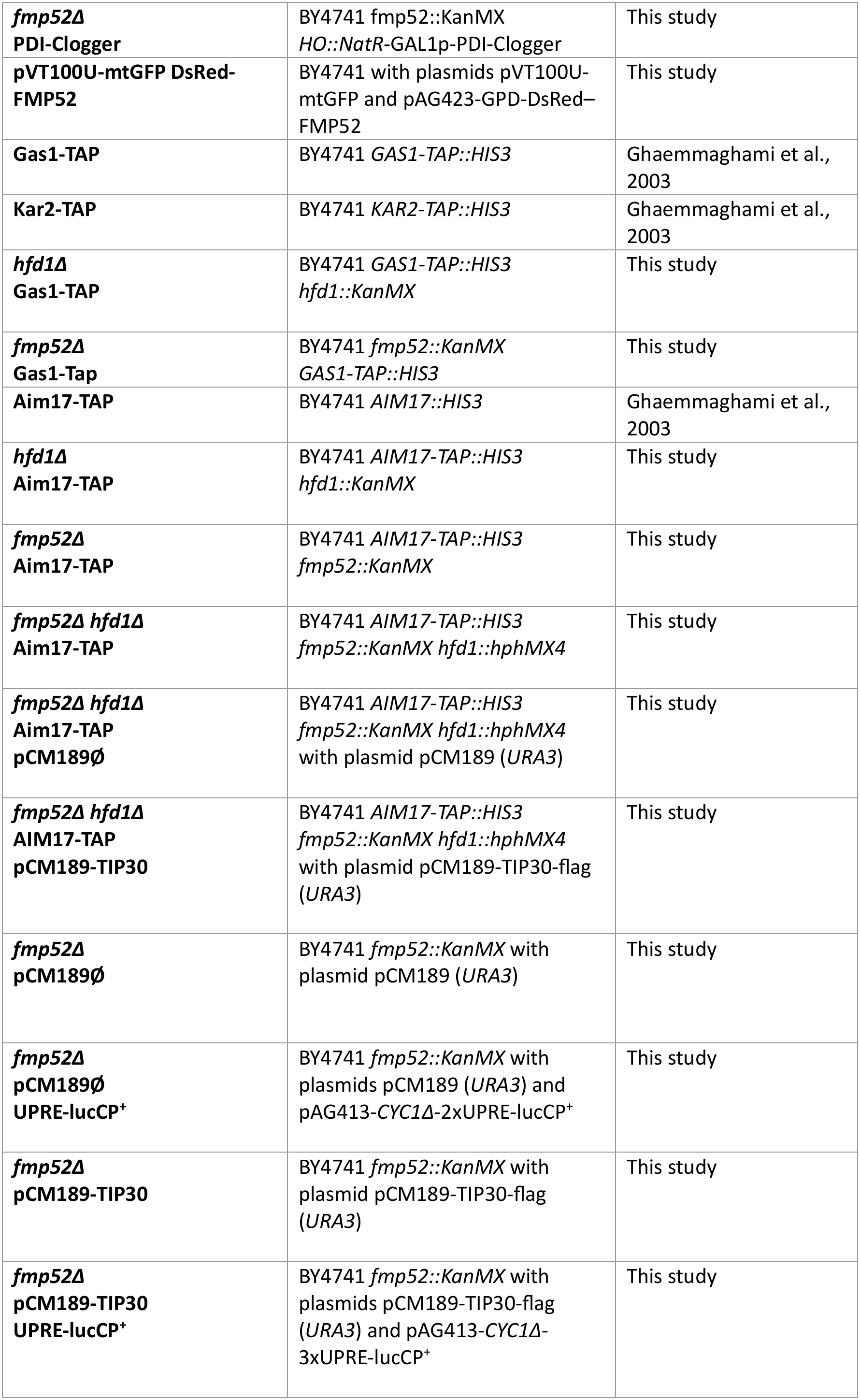
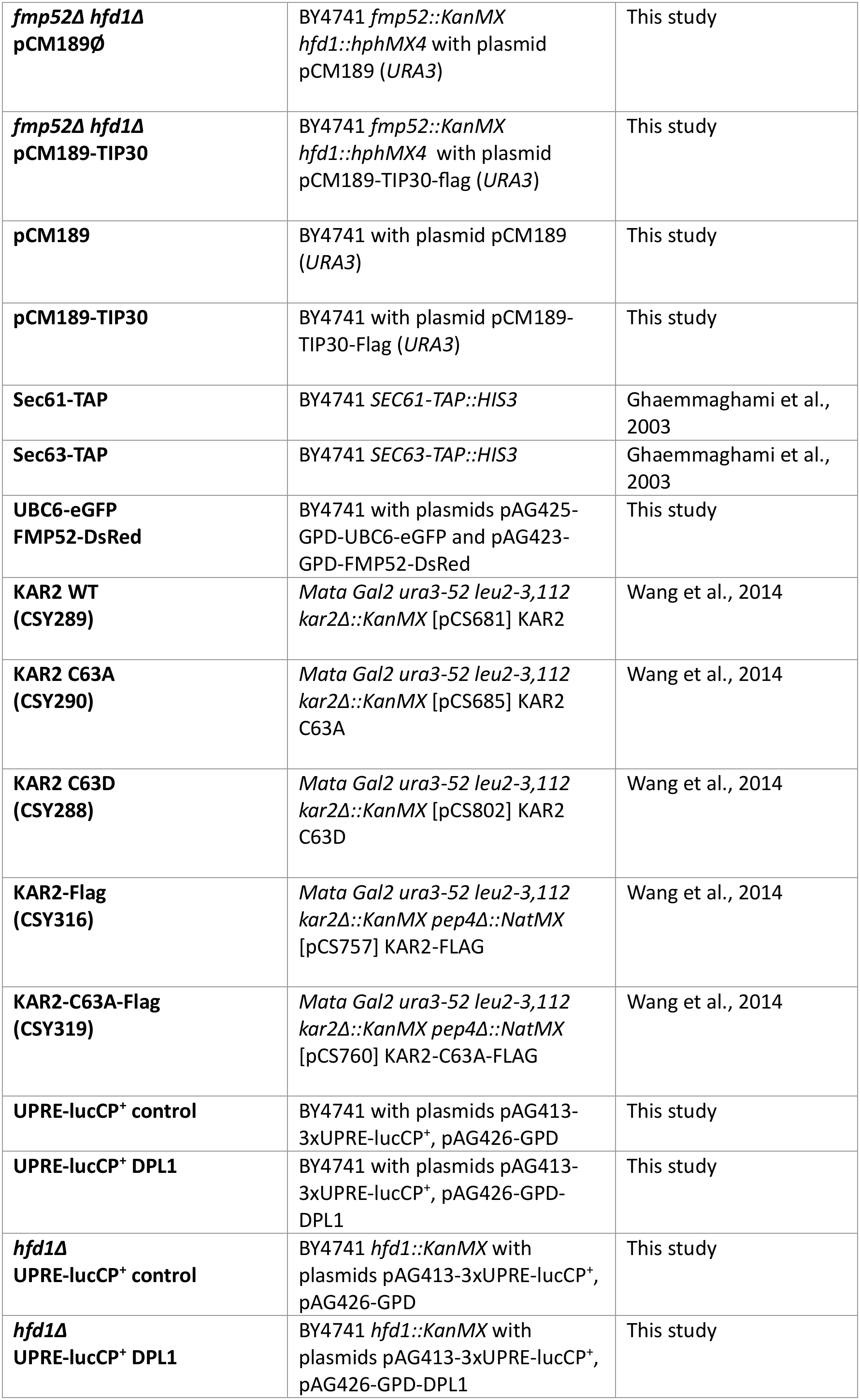
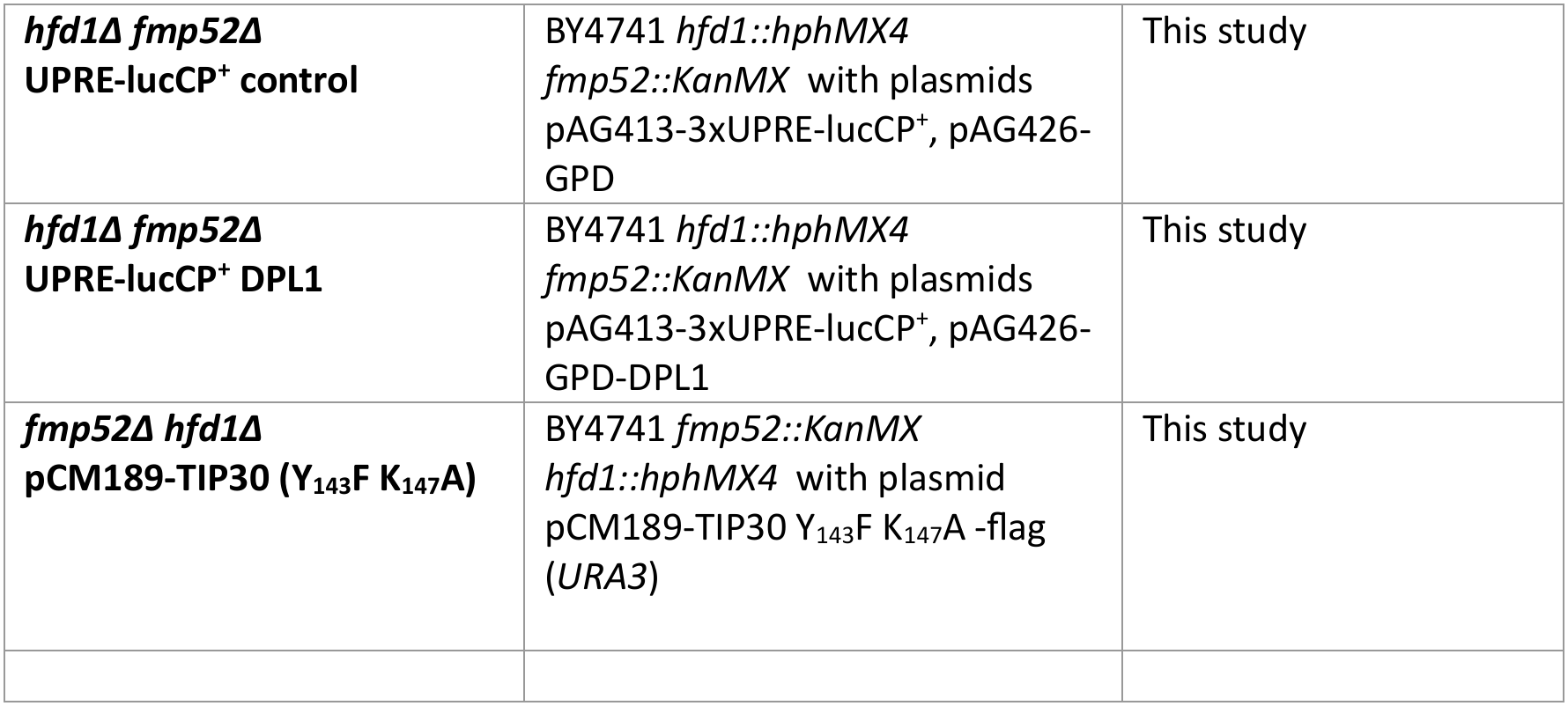
Yeast strains used in this study.

Bioactive lipids t-2-hex and t-2-hex-H_2_ were purchased from Merck and applied directly to yeast cultures from 1mg/ml stock solutions in DMSO.

The Kar2 C63A and C63D experiments were performed using previously established Kar2 replacement strains (CSY290 and CSY288, respectively) and the corresponding WT control strain (CSY289) described by Wang et al. ^31^. These strains contain a chromosomal kar2Δ::KanMX allele and are maintained by plasmids expressing WT KAR2, KAR2-C63A, or KAR2-C63D, respectively. These strains have a genetic background distinct from the BY4741-derived strains used for the majority of this study.

### Plasmids

The single- and multi-copy destabilized reporter plasmids pAG413-lucCP^+^ and pAG423-lucCP^+^ were described previously ^55^. The FMP52p-lucCP^+^ reporter was constructed by cloning the entire *FMP52* promoter (−693/-2) into the *Sac*I and *Sma*I sites of pAG423-lucCP^+^ ^55^. For the constitutive overexpression of *DPL1*, the plasmid pAG426-GPD-DPL1 was used ^15^. For the assay of specific cis-regulatory elements, synthetic oligonucleotides containing three repetitions of PACE ^19^ and of UPRE sequences were inserted into the *Bsp*EI site of plasmid pAG413-CYC1Δ-lucCP^+^ ^55^. The sequences used were the following (consensus sequence for each TF binding site is underlined): PACE, 5’-CCGG<u>CGGTGGCAAA</u>GATAT<u>CGGTGGCAAA</u>GTAAT<u>CGGTGGCAAA</u>T-3’; UPRE, 5’-CCGGATATCGA<u>CAGCGTG</u>TCATCGA<u>CAGCGTG</u>TCATCGA<u>CAGCGTG</u>TC-3’. For the visualization of the ER, the reporter plasmid pAG425-GPD-eGFP-UBC6 was used ^56^. A C-terminal fusion of Fmp52 with dsRed was obtained by cloning the entire *FMP52* ORF into the pAG423-GPD-ccdB-dsRed expression vector ^57^. For the expression of human Tip30 in yeast we cloned a synthetic DNA fragment (Twist Bioscience) comprising TIP30-Flag-6xHis into the Tet-off expression vector pCM189 ^58^. The TIP30 Y_143_F K_147_A mutant was generated by PCR-based site-directed mutagenesis using PfuUltra high-fidelity DNA polymerase and mutagenic primers carrying the desired nucleotide substitutions: TIP30-Y143F-K147A-fw, 5’-CAATTTTTTATTTCTACAAGTTGCGGGAGAAGTAGAAGCCAAG-3’and TIP30-Y143F-K147A-rev, 5’-CTTGGCTTCTACTTCTCCCGCAACTTGTAGAAATAAAAAATTG-3’. The complete plasmid (pCM189-TIP30-flag) was amplified for 12-18 cycles using an extension time of 1 min per kb of plasmid DNA. Following amplification, the reaction was treated with *Dpn*I for 1 h at 37°C to digest the methylated parental template DNA. *Dpn*I-treated DNA was subsequently transformed into competent *E. coli* DH5α cells, and plasmids carrying the desired mutations were identified and verified by DNA sequencing.

### Quantitative growth assays

For the quantitative estimation of growth parameters, fresh yeast overnight precultures of the indicated yeast strains were diluted in triplicate in the assay medium in multiwell plates to a starting OD_600_=0.1. Growth was then continuously monitored at a wavelength of 600nm every 30min at 28°C on a Tecan Spark multiplate reader for the indicated times and with two rounds of orbital shaking in between reads. The growth curves were processed in Microsoft Excel, and lag times and growth yields were calculated with the PRECOG software ^59^ with respect to control conditions.

### Quantitative cell death assays

Apoptotic cell death was quantified by terminal deoxynucleotidyl transferase dUTP nick-end labeling (TUNEL). Fresh yeast cultures were adjusted to an OD_600_ of 2 and treated with the indicated concentrations of trans-2-hexadecenal (t-2-hex) or solvent control for 4 h. Cells (1-2 × 10⁷) were harvested by centrifugation, washed twice with PBS, and fixed with 4% formaldehyde in PBS for 30 min at room temperature. After two washes with PBS, cells were resuspended in sorbitol buffer (1.2 M sorbitol, 0.5 mM MgCl₂, pH 8.0) and the cell wall was digested with 0.25 mg/mL Zymolyase 20T for 60 min at 37°C. Cells were subsequently permeabilized with 0.1% Triton X-100 in PBS for 2 min on ice and washed twice with PBS. TUNEL labeling was performed by incubating permeabilized cells with terminal deoxynucleotidyl transferase (TdT) and fluorescently labeled dUTP for 60 min at 37°C in the dark according to the manufactureŕs instructions (In Situ Cell Death Detection Kit Ref. 11684795910, Roche, Germany). Following two washes with PBS, TUNEL-positive cells were quantified by flow cytometry using a MACS Quant 10 flow cytometer (Miltenyi Biotec) using GFP/FITC fluorescence settings (488 nm excitation and 525/550 nm emission). 20,000 cells were analyzed for each measurement in triplicate, and aggregated cells (<5%) were excluded from further analysis.

Cell death was quantified by propidium iodide (PI) staining followed by flow cytometry. Yeast cells from overnight cultures were adjusted to an OD_600_ of 2 and incubated under the indicated treatment conditions for 4 h at 30°C with agitation. Cells were harvested by centrifugation, washed once with PBS, and resuspended in PBS. For PI staining, cells were incubated with PI (1:200 dilution from a 1 mg/mL stock solution) for 15 min at room temperature in the dark. Heat-killed cells (100°C for 20 min) and unstained cells were used as positive and negative controls, respectively. Samples were analyzed by flow cytometry using excitation at 488 nm and fluorescence detection at 655-730 nm. A total of 20,000 events were acquired per sample, aggregated cells were excluded from further analysis, and cell death was quantified as the percentage of PI-positive cells.

### Preparation of total protein extracts

Yeast whole cell extracts were generated using the boiling method of alkaline pretreated cells described by ^60^. A total of 2.5 OD_600_ were processed for each individual extract.

### Western blot

The following primary antibodies were used for the immunological detection of specific proteins on PVDF membranes: anti-Tap (CAB1001 rabbit polyclonal antibody, Invitrogen, 1:10.000), anti-HA (12CA5 mouse monoclonal antibody, Roche, 1:10.000), anti-Pgk1 (22C5D8 mouse monoclonal antibody, Invitrogen, 1:5.000), anti-Flag (M2 mouse monoclonal antibody, Sigma, 1:10.000). Immunoblots were ECL developed using the secondary antibodies anti-rabbit IgG HRP (Cytiva NA934, 1:10.000) and anti-mouse IgG HRP (Cytiva NA931, 1:10.000). Blots were visualized with an Image Quant LAS4000 system.

### RT-PCR detection of *HAC1* mRNA splicing

The *HAC1* mRNA splicing assays were performed following established protocols ^61^. Briefly, total RNA was isolated using the hot phenol extraction method and reverse-transcribed into cDNA using SuperScript™ III reverse transcriptase (Invitrogen) with oligo(dT) primers, according to the manufacturer’s instructions. *HAC1* cDNA was subsequently amplified by PCR using gene-specific primers: Hac1fw (5ʹ-CGCAATCGAACTTGGCTATCCCTACC-3ʹ) and Hac1rev (5ʹ-CCCACCAACAGCGATAATAACGAG-3ʹ).

### Real-time luciferase assays

Time-elapsed assays with destabilized firefly luciferase reporters in living yeast cells were performed essentially as described before ^55^. Yeast strains containing the indicated luciferase fusion genes on centromeric or multicopy plasmids were grown at 28°C in Synthetic Dextrose (SD) medium lacking histidine at pH=3 to low exponential phase. Culture aliquots were then incubated with 0.5mM luciferin (free acid, Synchem, Germany) on a roller at 28°C for 1h. The cells were then transferred in 135μl aliquots to white 96 well plates (Nunc) containing the indicated concentrations of stressor or solvent. The light emission was continuously recorded on a GloMax microplate luminometer (Promega) in three biological replicas. Data were processed with Microsoft Excel. Raw data were normalized for the number of cells in each individual assay and set to 1 for time point 0 as indicated. For the comparison of induction folds, luciferase activities are shown throughout this work as relative light units (R.L.U). For the comparison of induction sensitivities, the maximal luciferase activity was set to 100% for the most responsive stressor concentration.

### Fluorescence microscopy

Exponentially growing yeast cells on SD media were visualized on a Leica SP8 confocal microscope with a HCXPL APO CS2 63x objective or on a Zeiss LSM980 confocal microscope with a 63x objective and Airyscan detector. GFP was visualized with 488nm excitation and 509nm emission and dsRed with 545nm excitation and 572nm emission wavelengths.

### t-2-hex-alkyne lipidation assays

Lipidation assays *in vitro* with the t-2-hex-alkyne probe were essentially performed as described previously ^18^. Yeast cells expressing Kar2-, Sec61-, Sec63- or Spf1-Tap from their natural promoters or control cells were grown to exponential growth phase in Yeast Extract Peptone Glycerol/Ethanol (YPGE) medium containing 3% glycerol/2% ethanol. Cells were washed 1x with cold H_2_O and 1x with cold TSB buffer (10mM Tris/HCl pH 7.5, 0.6M sorbitol). Cells were lysed by mechanical rupture with 0.5mm glass beads in TSB buffer supplemented with 1mM PMSF and complete protease inhibitor cocktail (Roche). The mitochondrial/ER fraction was enriched from the lysates by two rounds of differential centrifugation: 5min at 3500g discarding the pellet and subsequent centrifugation of the supernatant at 16000g for 10min. The final pellet was resuspended in 250μl of HEPES lysis buffer (50mM HEPES pH 7.4, 0.5% NP-40) supplemented with complete protease inhibitor cocktail (Roche). Total protein was quantified with the Bradford method, and the mitochondrial fractions adjusted to 0.1mg/ml in Hepes lysis buffer. 500μl aliquots were used for *in vitro* lipidation experiments. The indicated concentrations of t-2-hex-alkyne ((E)-2-Hexadecenal Alkyne, 20714 Cayman Chemicals) or solvent alone were added from a 1mg/ml stock in ethanol and the samples were rotated at room temperature for 1h. After removal of the input samples (100μl), the rest of the samples was treated with the click reaction mix (40μl) of 100μM TBTA (678937 Sigma), 1mM TCEP (C4706 Sigma), 1mM CuSO_4_, 100μM Azide-PEG3-biotin conjugate (762024 Sigma) for 1h at room temperature. Samples were finally precipitated by adding 3.4ml chilled methanol over night at -80°C. Proteins were pelleted by centrifugation at 17000g for 15min and dried in a Speed Vac centrifuge for 5min. Proteins were first dissolved by addition of 50μl PBST (8mM Na_2_HPO_4_, 150mM NaCl, 2mM KH_2_PO_4_, 3mM KCl, 0.05% Tween 20, pH 7.4) containing 1% SDS and then diluted in 950μl PBST. Protein mixtures were incubated with 40μl PBST washed Streptavidin agarose resin (50% slurry, 69203-3 Merck) and incubated for 2h at room temperature on a roller. Agarose beads were washed 3x with PBST and proteins eluted with 80μl of elution buffer (95% formamide, 10mM EDTA) for 5min at 95°C. The supernatant was supplied with 1x Laemmli buffer and proteins were analyzed by immunoblotting with anti-Tap antibodies.

To determine whether Kar2 is modified in intact yeast cells, culture aliquots were incubated for 1h with the clickable lipid analogue t-2-hex-alkyne or solvent alone as a control, followed by cell lysis, CuAAC-mediated biotin labeling, and streptavidin enrichment as described for bioorthogonal alkyne-lipid probes with minor modifications ^62^. Following incubation, cells were extensively washed with HEPES lysis buffer (50 mM HEPES, pH 7.4, 0.5% NP-40) to remove excess probe before lysis. Cell extracts were then subjected to click chemistry and streptavidin purification under the same conditions used for the in vitro lipidation assays described above.

### Global analyses of t-2-hex transcriptional response, proteomic targets and genetic factors of tolerance

The transcriptomic response of yeast *hfd1Δ* mutants upon t-2-hex stress (100μM) was determined by us by RNA seq previously ^19^. All RNAseq data from this study are available to download from ArrayExpress (https://www.ebi.ac.uk/biostudies/arrayexpress/studies) under dataset accession number E-MTAB-13188.

Chemoproteomic profiling of t-2-hex targets using the lipid analogue t-2-hex-alkyne in mitochondria/ER-enriched yeast extracts has been reported in our previous work ^19^, where the complete proteomic dataset is available.

We previously performed a saturated transposon mutagenesis (SATAY) screen in a yeast *hfd1Δ* mutant to identify genes that positively or negatively influence t-2-hex tolerance ^19^. The complete SATAY dataset is available for download from ArrayExpress under accession number E-MTAB-13208.

### Statistical analyses

Data are presented as mean ± SEM unless otherwise indicated. The exact number of independent biological replicates (*n*) is specified in the corresponding figure legends. Comparisons between two groups were performed using two-sided unpaired Student’s *t*-tests. No formal tests for normality or equality of variance were performed because of the small sample sizes. Statistical analyses were performed using GraphPad Prism 8.0.1. A *p*-value < 0.05 was considered statistically significant.

## Acknowledgements

We would like to thank Susana Rodríguez-Navarro, Maya Schuldiner and Carolyn Sevier for the kind gift of yeast strains. We thank Rosa Viana for assistance and help with confocal microscopy image acquisition and Valentina Suarez for help with constructing luciferase reporters. We acknowledge support from the Scientific Network Enfermedades Metabólicas (COMETA) and Enfermedades Raras (RER-CSIC) funded by the Consejo Superior de Investigaciones Científicas (CSIC), Spain.

---

**Supplementary Figure 1.**
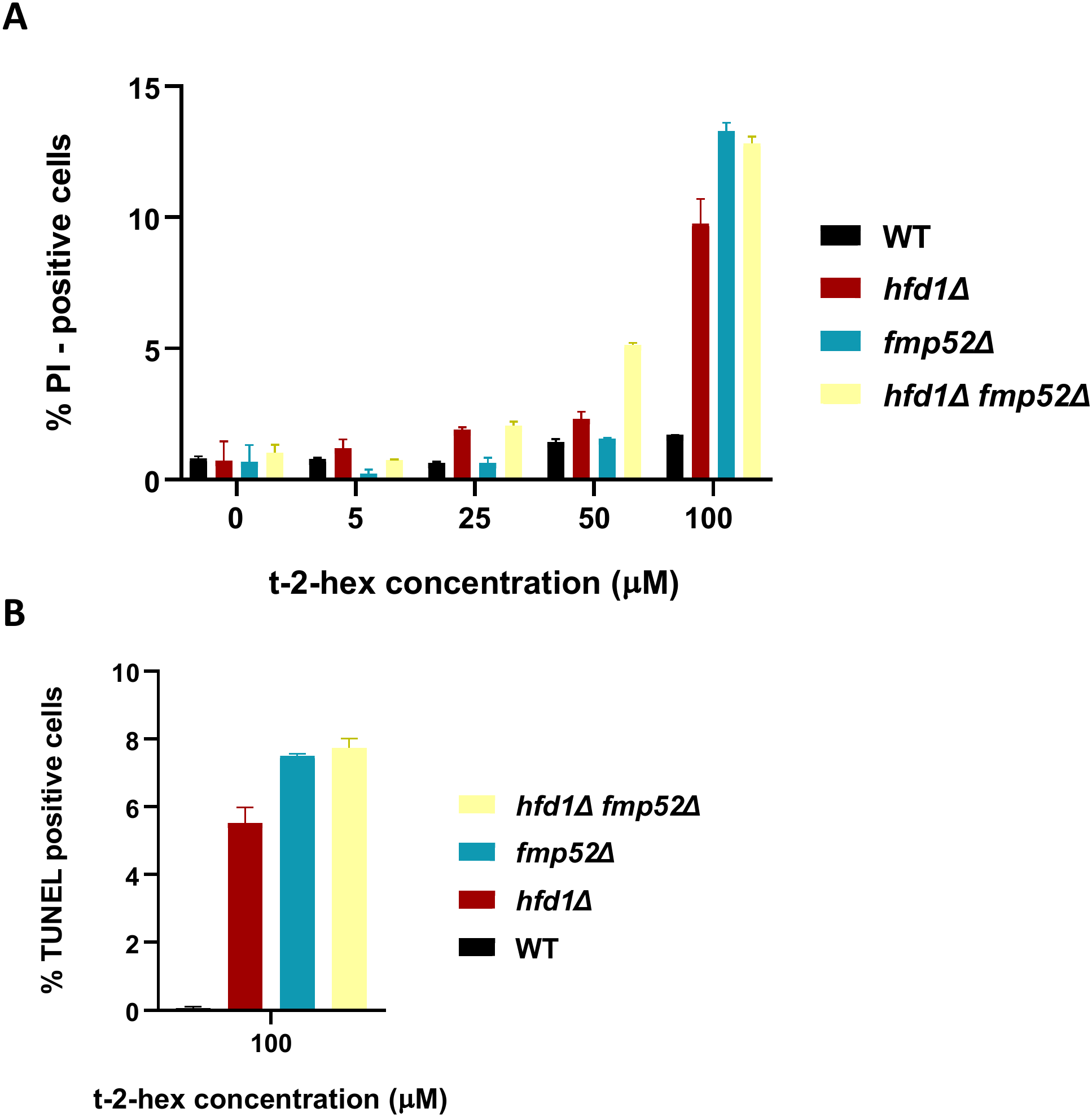
Quantification of general and apoptotic cell death upon t-2-hex exposure in wild type and detoxification mutants. (A) Quantification of propidium iodide (PI)-positive cells following t-2-hex exposure. The indicated yeast strains were treated with the indicated concentrations of t-2-hex for 4 h, and PI-positive cells were quantified by flow cytometry. (B) Quantification of apoptotic cell death by TUNEL assay. The indicated yeast strains were treated with 100 µM t-2-hex for 4 h, and TUNEL-positive cells were quantified by flow cytometry. Data are presented as mean ± SD from three biological replicates.

**Supplementary Figure 2.**
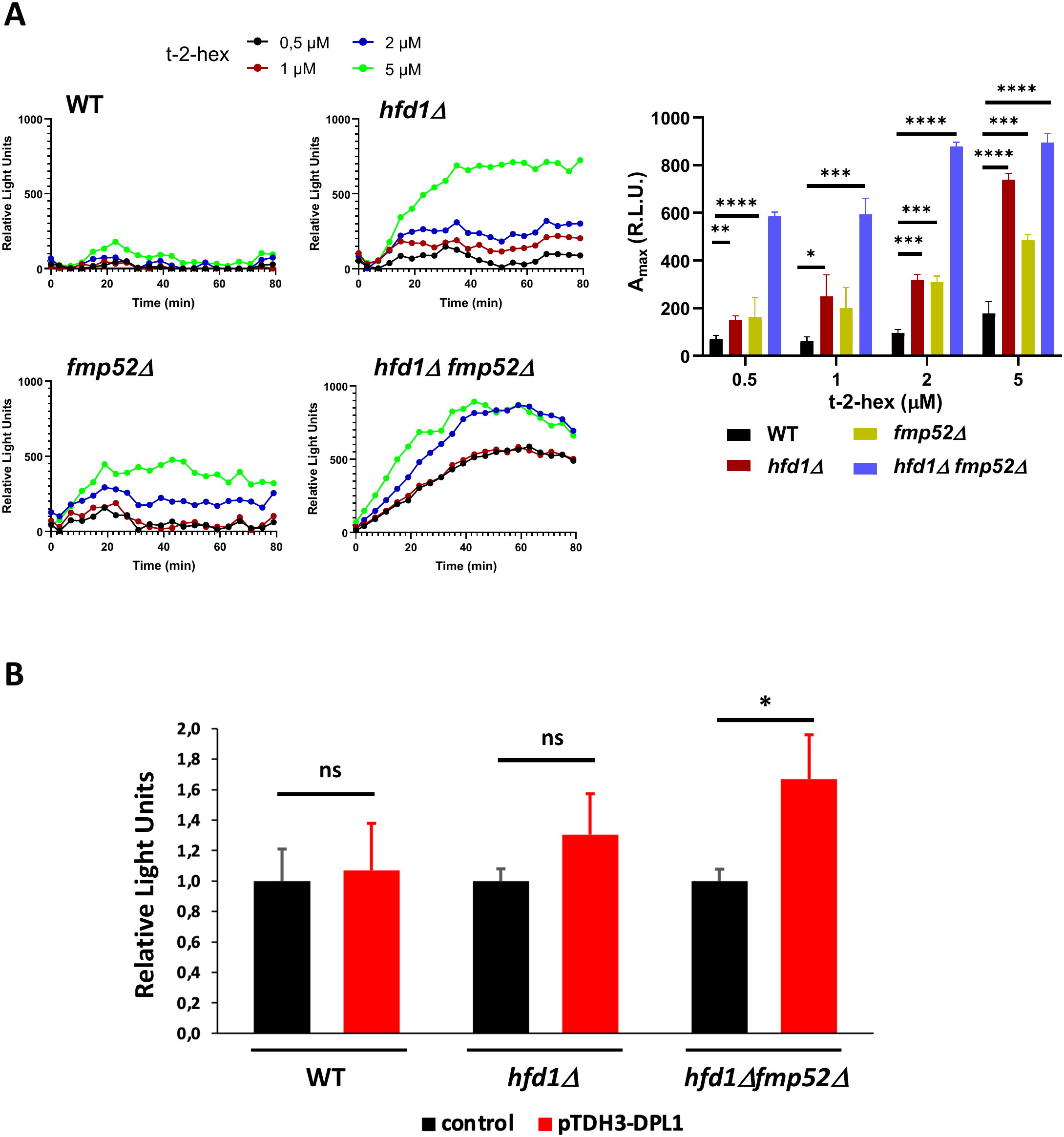
Low levels of exogenous t-2-hex and enhanced endogenous lipid production induce ER stress when lipid detoxification is impaired. (A) Quantitative live-cell luciferase assays using a UPRE-luciferase reporter in yeast wild-type, *hfd1Δ*, *fmp52Δ*, and *hfd1Δ fmp52Δ* cells exposed to the indicated low doses of t-2-hex. Data were normalized to the corresponding mock-treated controls. Left panels: Continuous dose-response curves. Right panel: Comparison of maximal UPRE activity at the indicated lipid doses. Data are mean ± SD from three biological replicates. (B) UPRE-luciferase activity in wild-type, *hfd1Δ*, and *hfd1Δ fmp52Δ* cells upon *DPL1* overexpression. Cells contained the UPRE-luciferase reporter together with either the *pTDH3-DPL1* overexpression plasmid or the corresponding empty vector as control. Data are mean ± SD from three biological replicates. *p<0.05, **p<0.01, ***p<0.001, ****p<0.0005 by Student’s unpaired t-test; ns, not significant.

**Supplementary Figure 3.**
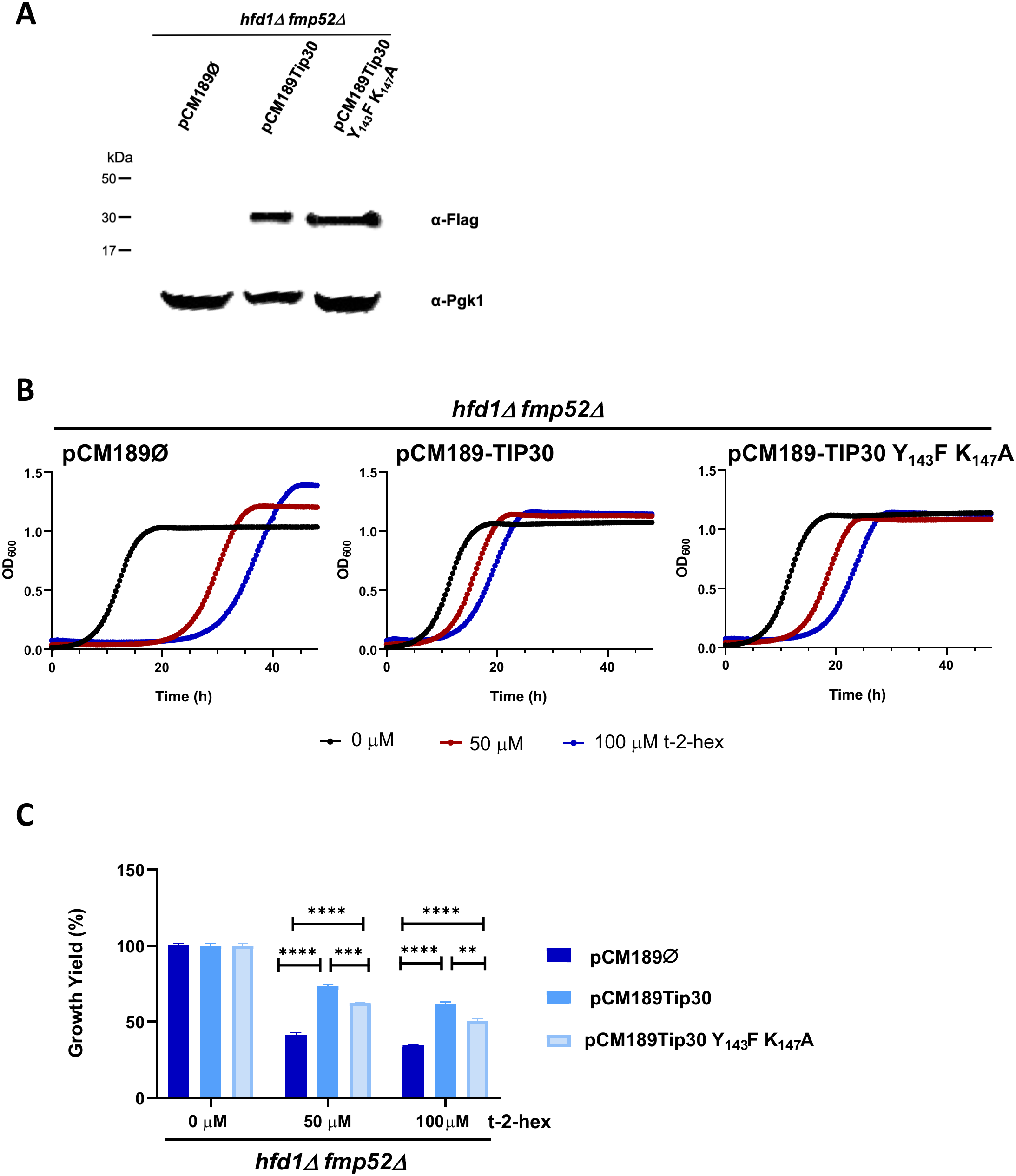
Functional analysis of the Tip30 Y_143_F K_147_A mutant targeting the putative catalytic YxxxK motif. Human Flag-tagged Tip30 variants were constitutively expressed in *hfd1Δ fmp52Δ* cells. (A) Expression of wild-type Tip30 and the Tip30 Y_143_F K_147_A mutant was analyzed in yeast total protein extracts by anti-Flag immunoblotting. Pgk1 was used as a loading control. (B) Quantitative growth assays of *hfd1Δ fmp52Δ* cells expressing wild-type human Tip30 or the Tip30 Y_143_F K_147_A mutant in the presence of the indicated t-2-hex concentrations. Shown are mean values from three biological replicates. (C) Quantification of t-2-hex dependent growth inhibition from the continuous growth experiments shown in (B). Data are mean ± SD from three biological replicates. **p<0.01, ***p<0.001, ****p<0.0005 by Student’s unpaired t-test.

